# Functional strain redundancy and persistent phage infection in Swiss hard cheese starter cultures

**DOI:** 10.1101/2021.01.14.426499

**Authors:** Vincent Somerville, Hélène Berthoud, Remo S. Schmidt, Hans- Peter Bachmann, Yi Hélène Meng, Pascal Fuchsmann, Ueli von Ah, Philipp Engel

**Author notes:** Corresponding authors, Department of Fundamental Microbiology, Batiment Biophore, University of Lausanne, 1015 Lausanne, Switzerland.

## Abstract

Undefined starter cultures are poorly characterized bacterial communities from environmental origin used in cheese making. They are phenotypically stable and have evolved through domestication by repeated propagation in closed and highly controlled environments over centuries. This makes them interesting for understanding eco-evolutionary dynamics governing microbial communities. While cheese starter cultures are known to be dominated by a few bacterial species, little is known about the composition, functional relevance, and temporal dynamics of strain-level diversity. Here, we applied shotgun metagenomics to an important Swiss cheese starter culture and analyzed historical and experimental samples reflecting 82 years of starter culture propagation. We found that the bacterial community is highly stable and dominated by only a few coexisting strains of *Streptococcus thermophilus* and *Lactobacillus delbrueckii subsp. lactis*. Genome sequencing, metabolomics analysis, and co-culturing experiments of 43 isolates show that these strains are functionally redundant, but differ tremendously in their phage resistance potential. Moreover, we identified two highly abundant *Streptococcus* phages that seem to stably coexist in the community without any negative impact on bacterial growth or strain persistence, and despite the presence of a large and diverse repertoire of matching CRISPR spacers. Our findings show that functionally equivalent strains can coexist in domesticated microbial communities and highlight an important role of bacteria-phage interactions that are different from kill-the-winner dynamics.

## Introduction

Natural microbial communities are complex biological systems. They are typically composed of a large number of different species and a high extent of strain-level diversity^1^. Moreover, they can be highly dynamic, influenced by microbial dispersion/migration, variation in physicochemical properties of the environment, nutrient availability, habitat size, interspecies interactions, and phage predation^2^. Therefore, it has remained challenging to track microbial communities in open biological systems and to understand how different factors shape their diversity, and eco-evolutionary dynamics.

Microbial communities harnessed by humans for the production of fermented foods are excellent models to investigate these questions because (i) they are less complex and (ii) are grown under much more stable conditions than present in nature^3^.

Cheese making relies on bacterial starter cultures, which have been propagated in cheese or milk over many generations, and are composed of a few coexisting species^4^. These starter cultures are essential for initiating the cheese making process by degrading proteins and fermenting lactose into lactate, resulting in the acidification and preservation of the milk environment. Two different types of cheese starter cultures (defined and undefined) are being used in today’s cheese-making industry^5^. Defined starter cultures are artificial communities that have been assembled from selected strains based on desired phenotypic properties^6^. On the contrary, undefined starter cultures are domesticated communities of unknown composition that were originally isolated from traditional cheese, but since then have been propagated and shaped by cycles of freeze drying, reactivation, and growth in milk. One of the major advantages of undefined starter cultures in cheese making is that they are phenotypically more stable and less prone to undergo spontaneous community collapse as frequently observed for defined cultures^7^. The underlying reasons have however not yet been elucidated. It is possible that diverse strains or species present in undefined communities makes them more resilient to environmental stressors^8^.

While the abiotic conditions usually remain highly stable when growing the starter culture in milk to produce cheese, bacteriophages are relatively common and pose a risk for the cheese making process^9^. Infection with a virulent phage can cause bacterial lysis, which in turn slows down fermentation and results in a low quality product, or the complete failure of the cheese production^10^. For a mesophilic undefined cheese starter culture, it was shown that different strains of *Lactococcus lactis* differ in their susceptibility to phages and that frequency-dependent negative selection, in the form of kill-the-winner bacteria-phage interactions, possibly prevent community collapse^11^. While it has been suggested that high levels of strain diversity may promote the functional stability of these communities, evolutionary theory predicts that strong bottlenecks such as those imposed during the starter culture propagation should purge diversity. We currently lack quantitative insights into the composition, functional relevance, and temporal dynamics of strain-level diversity in domesticated microbial communities^15^, and how it links to phage interactions.

Here, we focused on thermophilic starter cultures (RMK202) of a traditional Swiss hard cheese. Genomes of two dominant bacterial species, namely *Streptococcus salivarius subsp. thermophilus* (hereafter *S. thermophilus*) and *Lactobacillus delbrueckii subsp. lactis* (hereafter *L. delbrueckii*), have previously been sequenced, and the presence of several active phages and CRISPR-Cas defense systems identified indicating high phage pressure^16^.

We hypothesize that the microbial community in this starter culture is shaped by the repeated propagation cycles, and predict that the resulting population bottlenecks, in combination with the stable environment, caused a reduction of microbial complexity on multiple levels, i.e. species, strain, and phage diversity, as well as genomic content). To test this hypothesis we used longitudinal resolved shotgun metagenomics and analyzed the bacterial and viral diversity across multiple passages of this starter culture, including historical samples ranging from 1996 to 2019, as well as samples from an experimental propagation, simulating approximately 60 years of starter culture production. We then tested if the retained diversity contributes to important functional properties, such as flavor volatile diversity, positive species interaction and enhanced community resilience to e.g. phage invasion. To this end, we isolated 39 strains covering the large majority of strain diversity detected in the metagenomes and tested their functional properties based on metabolomic profiles, acidification potential, and growth dynamics. Finally, we extracted all CRISPR spacers and matched them to phages identified across the analyzed samples. Our results show that coexisting strains of the same species are functionally redundant, but differ tremendously in their phage resistance potential. Phages and bacteria seem to stably coexist over time, indicating a complex life cycle and suggesting functional implications for the cheese starter culture.

## Results

### Ongoing genome decay and putative species interaction based on metagenome-assembled-genomes

To obtain a reference for the genomic diversity present in the Swiss hard cheese starter culture RMK202, we applied Nanopore long read and Illumina short read sequencing to the starter culture from 2019. From this combined metagenomic data, we assembled two circular bacterial metagenome-assembled-genomes (MAGs) (Fig. 1A, Supplementary Fig. 1). These MAGs were most similar to *Streptococcus thermophilus* STH_CIRM_19 (ANI=99.5%,cov=94.1%) and *Lactobacillus delbrueckii subsp. lactis* KCTC 3034 (ANI: identity=99.1%,cov=82%), isolated from a French hard cheese Gruyère de Comté in 1963 and from sour milk in 1999, respectively. The two MAGs (hereafter referred to as Stherm_MAG and Lacto_MAG) are 1.9 Mb and 2.2 Mb in length, contain 1975 and 2163 genes, and six and nine complete copies of the rRNA operons, respectively. They both harbored a high number of pseudogenes (Stherm_MAG = 227, Lacto_MAG = 240) and many transposases (*S. thermophilus* = 75, *L. delbrueckii* = 212), which is in line with our hypothesis and previous reports suggesting ongoing genome decay as a result of microbial domestication in dairy communities^17,18^.

**Figure 1.**
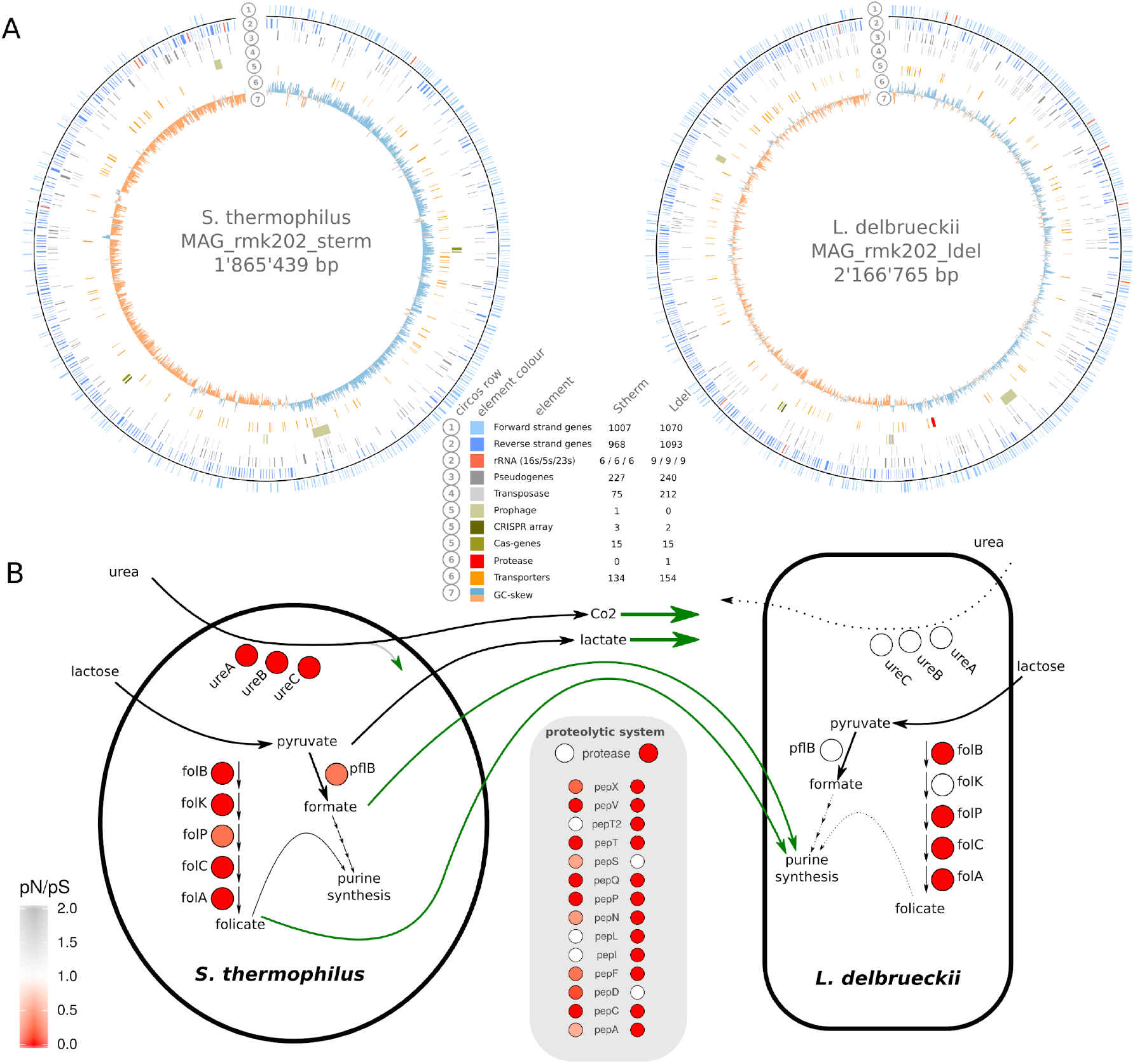
Assembly, annotation, and functional properties of the two metagenome-assembled genomes (MAGs) from the Swiss hard cheese starter culture RMK202. A) The Metagenome-assembled-genomes of *S. thermophilus* and *L. delbrueckii* with different genetic features highlighted (see legend). B) Functional properties potentially involved in the metabolic interaction of the two species. The coloring of the circles indicated the evolutionary conservation (pN/pS) of the genes, while empty circles indicate the absence.

We found a putative prophage sequence in the *S. thermophilus* MAG and multiple CRISPR arrays in both MAGs (Stherm_MAG = 3, Lacto_MAG = 2) indicating previous phage encounters. In addition, we were able to assemble three plasmids of which one had a very high and the other two a very low copy number (Supplementary Fig. 2). The three plasmids shared similarity to *Streptococcus suis* plasmid pISU2812 (id=95%), and *L. delbrueckii* plasmids pWS58 (id=95%) and pJBL2 (id=90%), but contained few genes with an annotation (Supplementary Fig. 2).

**Figure 2.**
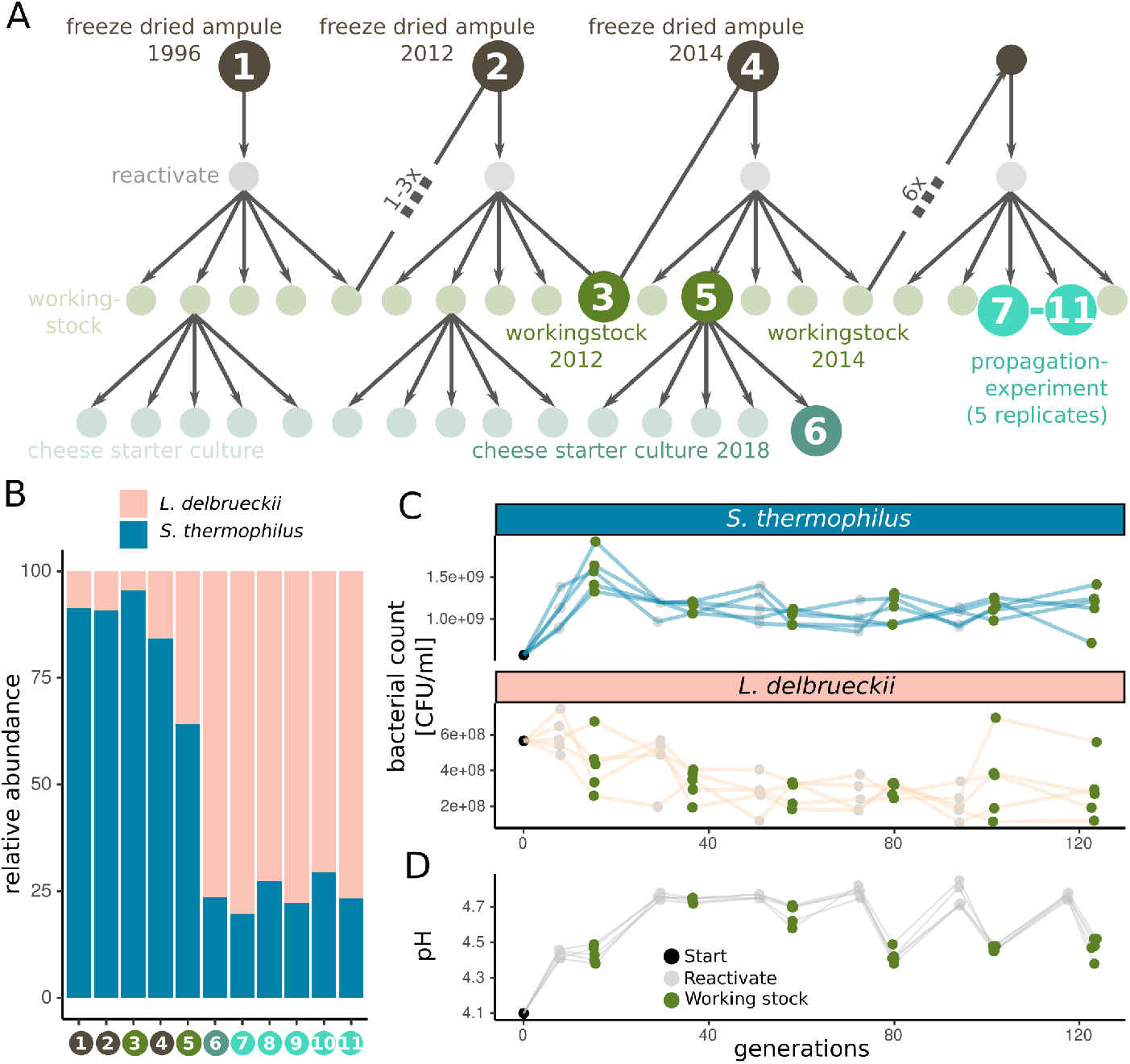
Metagenomic sampling design and species abundance. A) The starter culture propagation scheme as applied in the cheese starter culture production. The samples subjected to metagenomic sequencing are indicated by darker colors and labelled with numbers. Every propagation cycle includes a freeze drying, reactivation, and working stock step. From the working stock, commercial starter cultures for weekly shipments to cheesemakers are produced. The propagation experiment was carried out in the same way as in production and in five replicates corresponding to samples 7-11. The numbers between the working stock (x) indicate the number of cycles in between. B) The relative abundance of the two bacterial species in the eleven starter cultures samples (as illustrated in Fig. 2A). C) Bacterial counts throughout the propagation experiment for both species and the five replicates (lines are colored according to species and points according to samples within Fig. 2A). D) Acidification potential throughout the propagation experiment, as measured by pH reached after 18h incubation at 37 °C in milk.

In yogurt fermentation, *S. thermophilus* and *L. delbrueckii subsp. bulgaricus* have been described to engage in a mutualistic interaction by provisioning nutrients to each other^19,20^. Similar metabolic interactions may also occur between these two species in cheese starter cultures (Fig. 1B). Genes necessary for folate, lactate, and formate production were all present and under purifying selection in Stherm_MAG (pN/pS<1, Supplementary Fig. 3), while these pathways were partially or completely missing in Lacto_MAG (Fig. 1B). In turn, *L. delbrueckii* encoded the only casein-cleaving protease in the system (Fig. 1B). Proteases are assumed to be of crucial importance for peptide cleavage and flavour development in cheese production^21^. Together, these results suggest that both species are possibly maintained due to metabolic complementation and potentially positive effects on each other.

**Figure 3.**
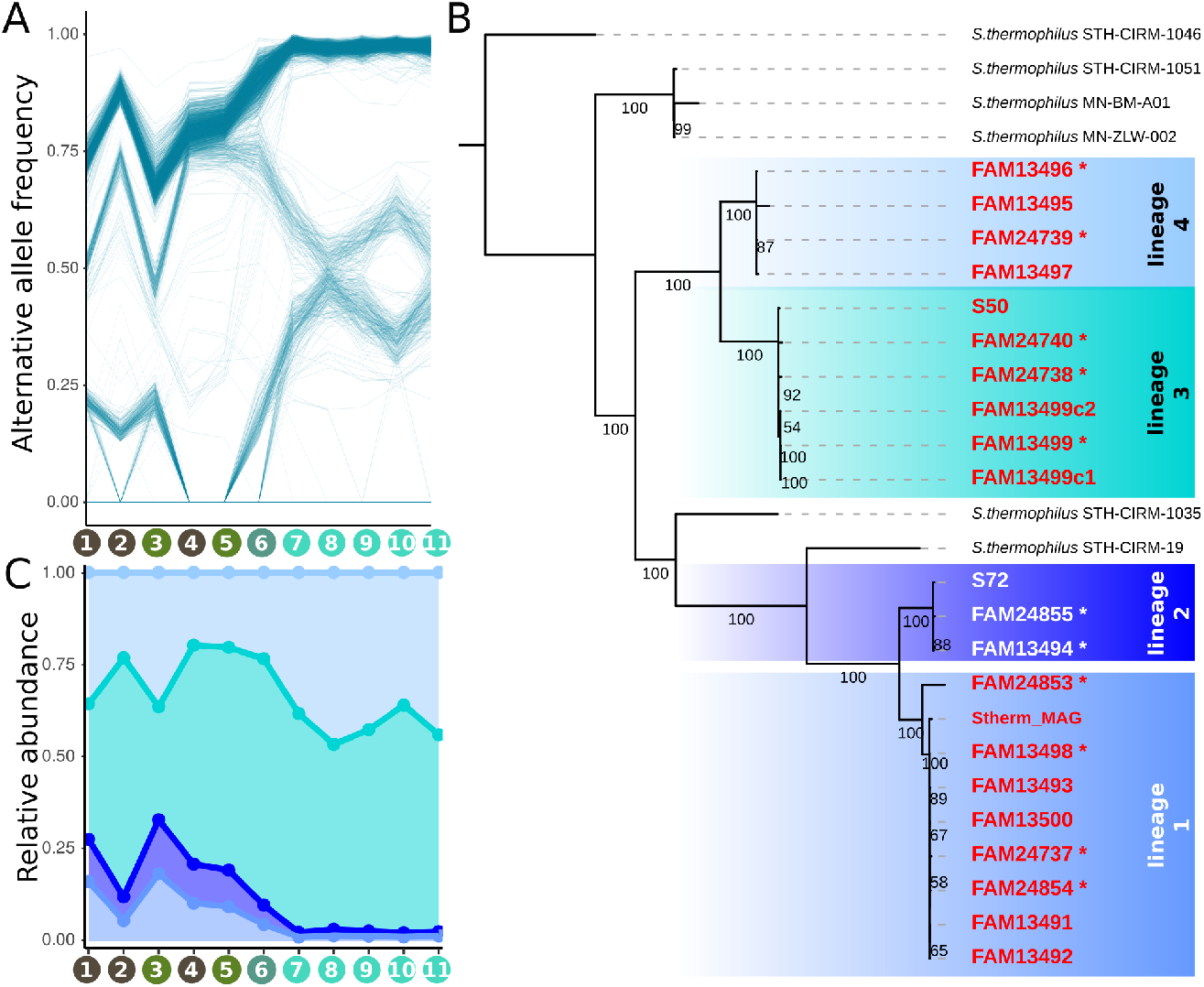
Strain-level diversity of *S. thermophilus* in cheese starter cultures. A) Alternative allele frequencies of all *S. thermophilus* SNVs over the metagenomic samples. Recurring SNVs from different samples are connected with a line. Clustering of lines indicates a large amount of SNVs with similar frequencies suggesting genomic coupling. Sample labels on the x-axis correspond to samples highlighted in Fig. 2A. B) The phylogeny of the isolated *S. thermophilus* strains based on maximum likelihood analysis on 1788 core genes. The isolates split into four lineages indicated by different color shadings. Strains sequenced with Nanopore are labelled with an asterisk. Values on branches indicate bootstrap values (100 replicates). C) The relative abundance of each of the four sub-lineages of *S. thermophilus* across the eleven metagenomes as based on the average frequency of lineage-specific SNVs identified on the basis of the isolates in Fig. 3B.

### Temporal stability of a two-species community in Swiss hard cheese starter cultures

To assess the effect of continuous passaging on the starter culture’s community structure and population dynamics, we documented the temporal dynamics of the community members. Cheese starter cultures are maintained by a systematic propagation scheme in order to minimise compositional shifts (Fig. 2A). In addition to the already characterized starter culture from 2019, we analysed six historical samples from 1996 to 2018 (9-15 propagation steps). Further, we continued to propagate the starter culture 12 times in five replicates, which is equivalent to 60 years of starter culture maintenance, or approximately 123 bacterial generations, considering a survival rate of 71.5% after freeze-drying (Fig. 2A, Supplementary Fig. 4). All samples were Illumina sequenced to a depth of 5-18 mio reads. We did not identify any additional community members, apart from the two species characterized in Fig. 1. To estimate the relative abundance of the two species, we mapped the metagenomic reads against the two MAGs (Fig. 2B). The historical samples revealed more variation in relative abundance of the two community members than the five replicates of the propagation experiment, most likely due to inconsistent sampling rather than real community differences. Colony counts of both *S. thermophilus* (mean=1*10^8^, SD=2*10^8^) and, SD=2*10^8^) and *L. delbrueckii* (mean=3*10^8^, SD=1*10^8^) remained similar throughout the 12 propagations (Fig. 2C), and all samples stably acidified to an average pH of 4.6±0.2, (Fig. 2D). The stable metabolic activity of both community members is further supported by the consistently high acidification rates (Supplementary Fig. 5) and constant ratios of D-lactate (i.e. *Streptococcus*) to L-lactate (i.e. *Lactococcus*) (mean=30%±10%; Supplementary Fig. 6) in the culture production plant since 1996. Together, these findings suggest that the two species stably coexist in the hard cheese starter cultures with little variation in abundance or phenotypic properties across time.

**Figure 4.**
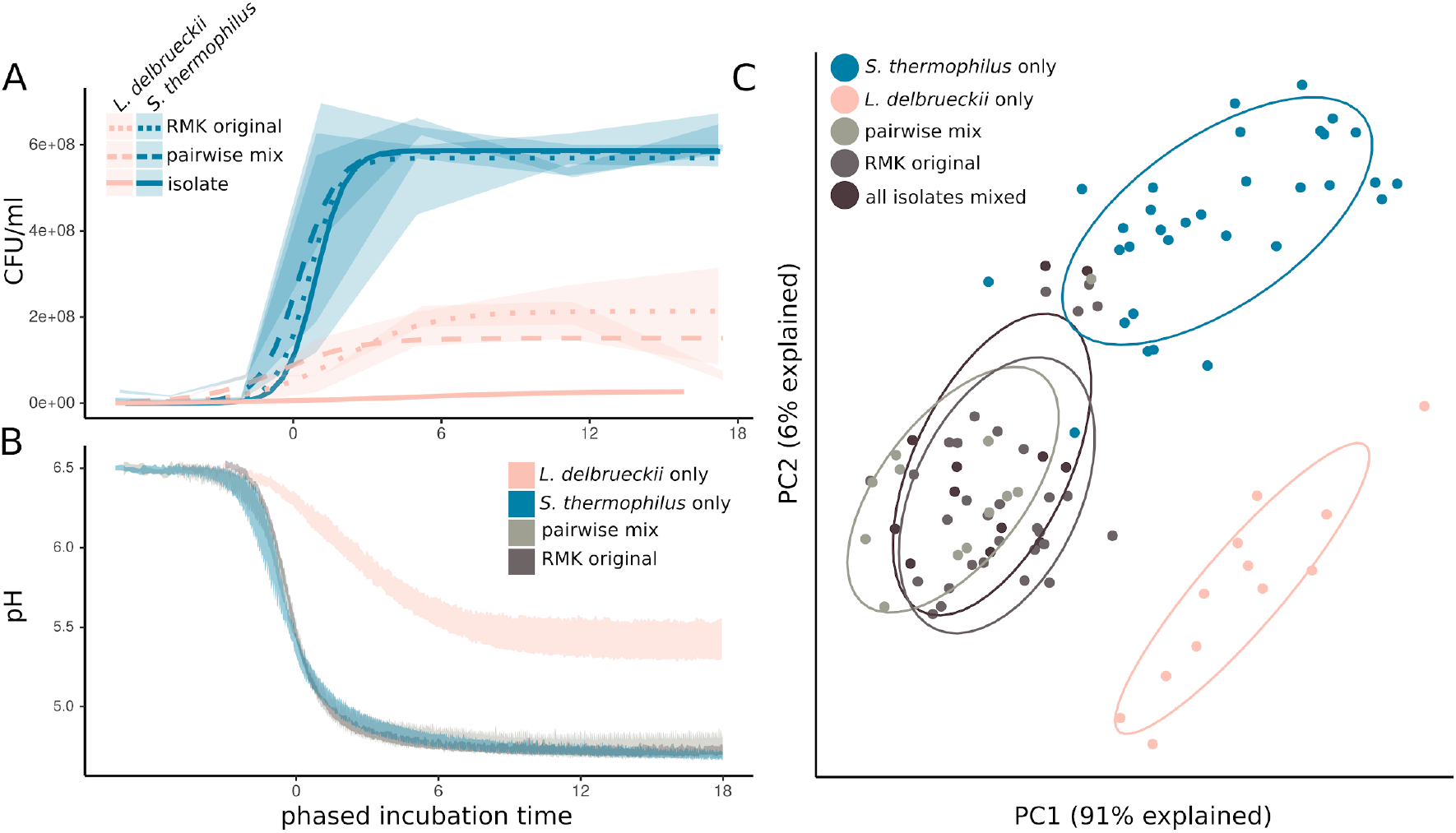
Phenotypic properties of individual strains, pairwise combination of strains, and original starter culture. A) Colony forming units (CFUs) of *S. thermophilus* and *L. delbrueckii* over 18h of growth when culturedalone, in pairwise combinations, or in the original starter cultures (RMK). The ribbons illustrate the interquartile rangeand the lines the modeled growth curves. B) Acidification curves of the same samples. The ribbons illustrate the min and max pH of the different samples. C) Principal component analysis of the metabolic profiles after 24 h of growth at 37 °C. Different treatments are highlighted in colors and with the surrounding eclipse.

**Figure 5.**
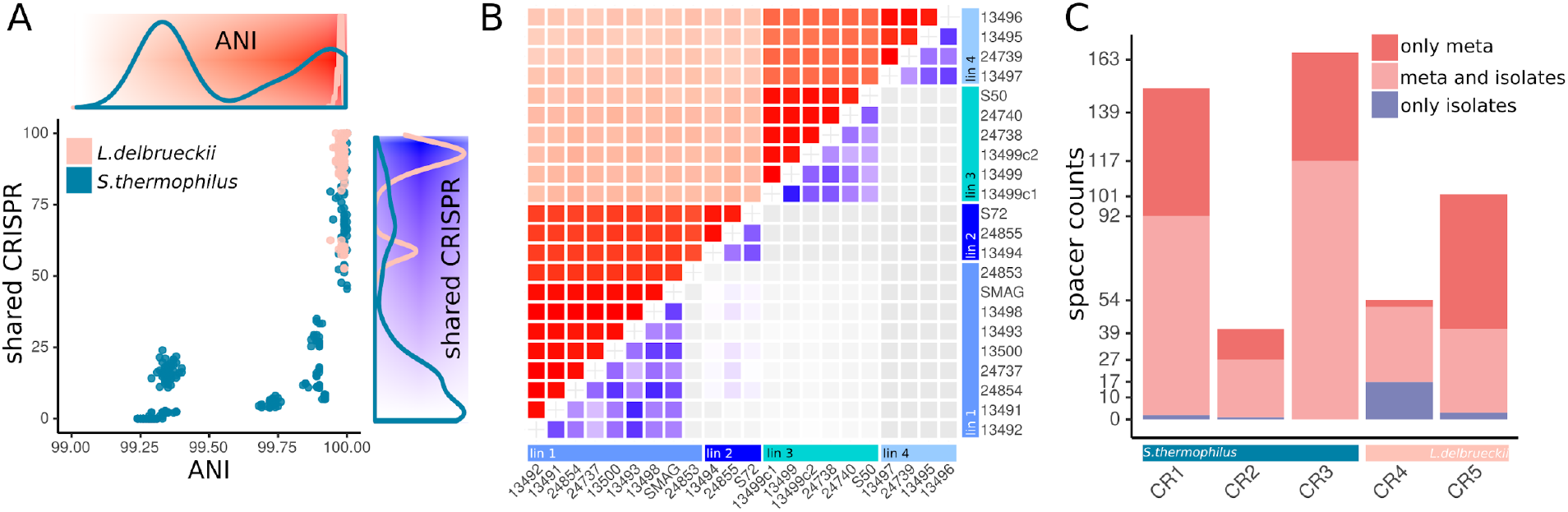
CRISPR spacer diversity of L. delbrueckii and S. thermophilus. A) The correlation of fraction of shared CRISPR spacers and ANI of all L. delbrueckii and S. thermophilus with the corresponding densities and heatmaps on the x and y-axis. B) The heatmap of the genomic and CRISPR spacer diversities of S. thermophilus illustrated with ANI (top heatmap; from white to red) and percent shared CRISPR spacers (bottom heatmap; from white to blue) C) The amount of metagenomic and genomic CRISPR spacers according to the five arrays.

**Figure 6.**
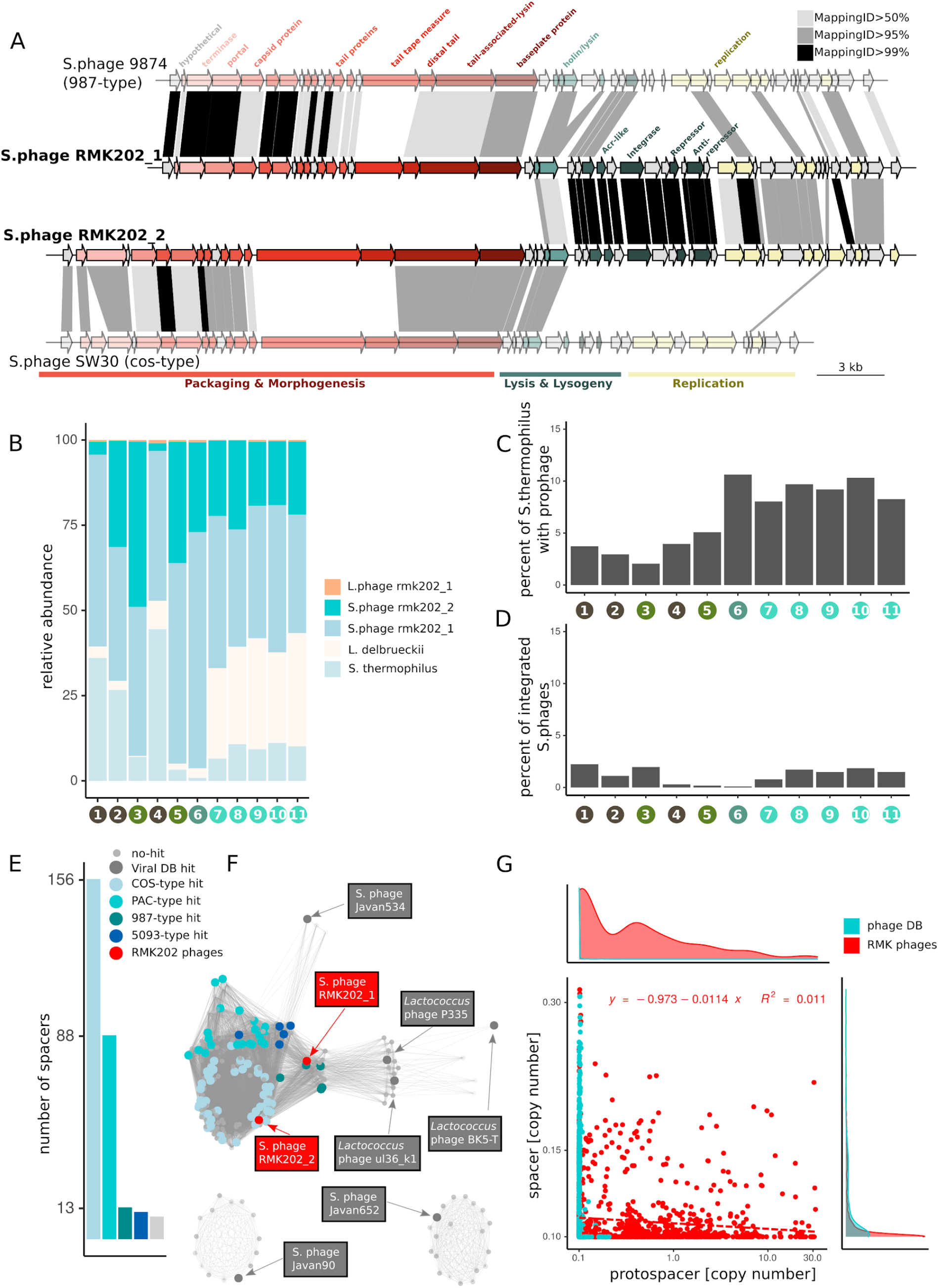
Characteristics of the phages identified in the cheese starter cultures. A) Gene annotation of the two *Streptococcus* starter culture phages, RMK202_1 and RMK202_2, and the two closest relatives (illustrated in lighter colors). Protein similarity between genes are indicated in grey (80-95% identity) and black (95-100%). B) Relative abundance of bacteria and phages over all metagenomic samples based on genome read coverage. C) Fraction of *Streptococcus* genomes with an integrated phage as based on the read coverage of phage-bacteria spanning regions relative to the coverage of the *S. thermophilus* genome. D) Fraction of *Streptococcus* phages which show signs of integration as based on the read coverage of phage-bacteria spanning regions relative to the coverage of the *Streptococcus* phage genomes. E) The number of spacers mapping against the different phage types. F) The *Streptococcus* phage network with the protospacer containing phages colored or labeled according to phage type. G) The spacer abundance versus the protospacer abundance from all phage spacers. The database specific linear regression and distributions are indicated in the figure and the axis figures accordingly.

### Low levels of intra-species diversity with a few stably coexisting *S. thermophilus* strains

While the starter culture community consists of only two stably coexisting species, there might be further variation at the strain-level. To assess the extent of intra-species diversity in the analyzed cheese starter cultures, we quantified single nucleotide variants (SNVs) across the core genes of the two community members. The fraction of variable sites in the 11 metagenomic samples differed between the two species. We found an average of 0.4% ± 0.002% (i.e. 4136 SNVs) variable sites for *S. thermophilus* and 0.0005% ± 0.0002% (i.e. 130 core genome SNVs) variable sites for *L. delbrueckii* (Supplementary Fig. 7). This is much lower than what has been reported for natural communities such as the human gut microbiome (∼4%)^22^ or the bee gut microbiome (∼10%)^23^ seems to confirm that the repeated propagation in a stable environment and the existence of strong population bottlenecks during freezing-propagation cycles reduced the strain-level diversity.

To identify co-existing strains of *S. thermophilus*, we looked at SNV frequencies across the 11 metagenomic samples. SNVs with similar frequencies in several independent samples are likely to be physically located on the same genomic element and consequently are contained in the same strains^24^. We found that the SNVs clustered into three (Supplementary Fig. 8) discrete phases (i.e. putative strains, Fig. 3A) which were consistently present across the eleven metagenomic samples. Only one phase transiently disappeared in two consecutive samples (Fig. 3A, sample 4 and 5), indicating that it fell below the detection limit. This analysis was not conducted for *L. delbrueckii*, given the low number of SNVs in this species.

To further characterize the variation within each of the two species, we cultured bacterial strains from the 2018 starter culture (sample 6). We genotyped the isolates using a mini-satellite region and sequenced their genomes using Illumina and Nanopore sequencing. This resulted in 22 and 17 draft or fully closed genomes of *S. thermophilus* and *L. delbrueckii*, respectively (Supplementary Table 1). Overall, the assembled genomes explained 97% core genome SNVs detected across the 11 metagenomes. The unexplained SNVs did not belong to any of the identified phases and were of lower abundance (Supplementary Fig. 9), indicating that we have cultured nearly the complete microbial diversity of the analyzed cheese starter cultures.

The isolated *L. delbrueckii* strains were highly similar to each other, containing 1732 core genes with the same 130 SNVs that were also identified across the eleven metagenomes (Supplementary Fig. 10). Only 118 genes were accessory, most of which had no annotation (n=53) or encoded transposases (n=59). Transposons are generally very common in *Lactobacillus* genomes (mean=163, sd=56) and are hard to assemble^25^.

On the contrary, the genomes of the *S. thermophilus* isolates were less similar to each other. The core genome consisted of 1788 genes with 4116 SNVs. The core genome phylogeny revealed that the *S. thermophilus* isolates clustered into four well-supported lineages, which were not monophyletic but separated by two French hard cheese (Comté) isolates (Fig. 3B). *S. thermophilus* harbored slightly more accessory genes (354) which were mostly annotated, similar as in *L*.*delbrueckii*, as hypothetical proteins (41%) or transposases (24%). A few accessory genes (n=21, i.e. 6%), however, encoded functions related to amino acid transport and polysaccharide biosynthesis (Supplementary Fig. 11), including eight glycosyl- and sugar transferase genes involved in cell wall modifications of *S. thermophilus* and known for their role in phage resistance^26,27^. The genomes from different lineages were largely syntenic (Supplementary Fig. 12). This explains, together with the relatively high similarity among the strains, why the MAG represented a single circular and non-chimeric genome (Fig. 1A).

Based on the isolated strains, we determined SNVs specific to each of the four lineages, and used their average frequency to estimate the abundance of each lineage across the metagenomic samples (Fig. 3C). This revealed that all four lineages stably coexist, but that, in particular within the propagation experiment, the lineage 3 and 4 dominate in the cheese starter cultures.

### Inter-species interactions and phenotypic redundancy among strains of the cheese starter cultures

While the composition of the community is simple, the two species and the multiple coexisting strains may contribute important functional properties. We experimentally assessed the functional differences across species and strains by looking at three important phenotypic properties of cheese starter cultures, namely the growth rate, acidification rate and the flavour volatiles production. We not only tested each isolate separately, but also included different pairwise strain combinations, a mix of all strains, and the original starter cultures.

For both species, all tested strains showed highly similar growth and acidification rates in mono-cultures. *S. thermophilus* isolates grew and acidified rapidly reaching a carrying capacity of 5.9*10^8^ CFU/ml, and decreasing the pH to 4.7, already 3h after the lag phase (Fig. 4A,B). In contrast, *L. delbrueckii* isolates grew and acidified significantly slower reaching a carrying capacity of 2.8*10^7^ CFU/ml, and decreasing the pH to 5.5, only 7h after the lag phase (Fig. 4A,B). Notably, culturing the two species together resulted in a 10-fold increase in the amount of *L. delbrueckii* from 2.8*107 CFU/ml to 2.2*10^8^ CFU/ml, in all strain combinations as well as in the original starter culture, suggesting a strong positive effect of *S. thermophilus* on *L. delbrueckii* growth. In contrast, neither were the cell counts of *S. thermophilus* higher in any of the co-culture conditions in comparison to the axenic cultures, nor did we observe a difference in acidification.

To identify potential differences in the formation of flavour volatiles among the individual strains, as well as the co-cultures, we compared the metabolic profiles after 24h of growth in milk. Principal component analysis identified three distinct clusters, two encompassing all strains of each species when grown alone, and one representing all co-cultures. This suggests that strains of the same species produce similar metabolic profiles, and that the metabolic profile changes when the two species are grown together, independent of how many and which strains were cultured together.

Taken together, these results suggest that interbacterial interactions between the two species modulate the flavour volatile profiles of starter cultures, but that any pairwise combination of strains reconstitutes the acidification properties and volatile profile of the starter cultures.

Hereby we reject our hypothesis that strain-level diversity contributes to flavor volatile diversity and conclude that the strains are functionally redundant.

### Highly variable phage resistance mechanisms in isolated genomes

Ecological theory predicts^28^ that functional redundancy prevents stable coexistence, which is in stark contrast with our results. We hypothesize that phage predation may contribute to the maintenance of low levels of strain diversity in the analyzed cheese starter culture, similar as suggested for mesophilic cheese starter cultures^11^. We can infer such interactions from genomic data, as bacteria store a memory of past phage encounters in their genome by integrating short spacer sequences into CRISPR arrays^29^. We found three previously described CRISPR-regions (CR) in the analyzed *Streptococcus* genomes, namely CR1 (type IIA), CR2 (type IIIA) and CR3 (type IIA)^30,31^ and two in *Lactobacillus*, CR4 (type IC) and CR5 (type III-A)^32^ (Supplementary Fig. 13). Despite little genetic divergence and accessory gene content variation, we found high variability in the number and identity of the CRISPR spacers in both species. A total of 236 and 92 unique CRISPR spacers were identified for *S. thermophilus* and *L. delbrueckii*, respectively. A given strain carried between 23 and 83 spacers (Supplementary Fig. 14), 75 spacers were only present in a single strain (Supplementary Fig. 15), and not a single spacer was shared by all *S. thermophilus* strains (Supplementary Fig. 16). Interestingly, the fraction of shared spacers between two genomes rapidly decreased with increasing genomic distance (Fig. 5A). Moreover, *S. thermophilus* strains of the same lineage shared between 58%-76% of their spacers, which is surprisingly little considering that the genomes within a lineage were nearly identical based on the ANI (ANI=99.999%, Fig. 5B). *S. thermophilus* CRISPR spacers present in only one strain tended to lie closer to the leader (Supplementary Fig. 17), where new spacers are usually acquired^33^.

To get a better understanding of the temporal dynamics of CRISPR spacer turnover, we extracted all reads containing CRISPR spacer sequences from the metagenomes, and assigned them to one of the five CRISPR arrays of *Lactobacillus* and *Streptococcus*, based on their conserved repeat sequence motifs (Supplementary Fig. 13). We identified a total of 354 and 131 dereplicated metagenomic CRISPR spacers for *S. thermophilus and L. delbrueckii*, respectively. Despite culturing the large majority of strain-level diversity (97%), 121 (34 %) and 64 (49%) spacers were not present in any of the isolated genomes of *S. thermophilus and L. delbrueckii* (Fig. 5C). Moreover, 15 *S. thermophilus* and 38 *L. delbrueckii* spacers were only found in the metagenomes of the propagation experiment. From these newly acquired spacers, and the known bacterial generation time, we estimated a mean CRISPR spacer turnover rate of 0.024 (SD=0.023) and 0.086 (SD=0.047) spacers per bacterial generation for *S. thermophilus* and *L. delbrueckii*, respectively. This is much lower than observed in previous experiments where *S. thermophilus* was challenged with new phages^34^. Together, these results suggest a continuous turnover of CRISPR spacers and ongoing bacteria-phage interactions.

### Two abundant and persistent *Streptococcus* phages in the cheese starter cultures

Our CRISPR spacer analysis suggests high rates of CRISPR spacer turnover and continuously high phage pressure. In order to characterize the viral community in the cheese starter cultures, we assembled all metagenomic reads that did not map against the bacterial MAGs (2-29%, Supplementary Fig. 18). A total of 23 non-redundant contigs were assembled on the basis of which we could reconstruct three complete phage genomes (rmk202_1-3, Supplementary Fig. 19). Adding these phages to our bacterial reference genomes (MAGs and plasmids, see above) resulted in the mapping of the large majority of all metagenomic reads (99.97%, SD=0.02%, Supplementary Fig. 20). Comparison with the viral RefSeq database indicated that phages rmk202_1 and rmk202_2 clustered with cos-type and 987-type phages of *Streptococcus* (Fig. 6A, Supplementary Fig. 21), respectively, whereas phage rmk202_3 clustered with the *Lactobacillus* phage JCL1032 (Supplementary Fig. 22).

Metagenomic read recruitment revealed that both *Streptococcus* phages were highly abundant, markedly surpassing bacterial numbers in several of the analyzed cheese starter cultures (1-110 phage copies per bacteria; Fig. 5C, Supplementary Fig. 23). In contrast, the *Lactobacillus* phage was much less abundant with an average of 0.1 phage copies per bacteria (SD=0.1x). Both *Streptococcus* phages showed little genetic variation: we only detected six mutations across the eleven samples despite the fact that both phages were sequenced very deeply (Supplementary Fig. 24). Remarkably, despite being highly divergent in most of the genome, the two *Streptococcus* phages shared a high degree (>99% nucleotide identity) of similarity in the lysis and lysogeny gene cluster coding for holin, lysin, integrase, repressor and anti-repressor suggesting a recent recombination event and a similar temperate lifestyle (Fig. 6A)^35–37^. The phage rmk202_2 was found to be integrated in the genome of only one of the 22 characterized *S. thermophilus* strains (FAM24739, Supplementary Table 1). Moreover, we identified metagenomic read pairs that mapped with one mate to the phage and the other to the *S. thermophilus* genome, providing further evidence for genomic integration. Based on the location of the mapped phage-bacteria spanning read pairs, we were able to identify four alternative integration sites (Supplementary Fig. 25). Moreover, a subset of the read pairs unambiguously mapped to each of the four lineages indicating that all lineages serve as host for the *Streptococcus* phages (Supplementary Fig. 26). Based on the relative number of such phage-bacteria spanning read pairs, we estimated that between 2-11% of all *S. thermophilus* cells carry an integrated phage (Fig. 6C), but that only between 0.1-2% of all *Streptococcus* phage genome copies are integrated (Fig. 6D). This highlights that the phages are active in the starter cultures and likely produce large numbers of phage particles.

Surprisingly, 35% of all metagenomic spacers identified above matched to one of the two *Streptococcus* phages. Moreover, each *Streptococcus* isolate, including the prophage-containing isolate, contained 3 to 15 spacers against the two phages (Supplementary Fig. 27). The presence of multiple spacers targeting the same phage suggests repeated encounters of that phage, as the integration of additional spacers is facilitated during CRISPR-mediated phage interference (known as interference-driven spacer acquisition^38^). The matching spacers were located throughout the CRISPR arrays (Supplementary Fig. 28), suggesting both relatively old as well as more recent acquisition events. Based on the above estimated spacer turnover rate (0.024 spacers/generation), and taking into account the distance of the spacer from the leader, we estimated that the first spacers targeting the two phages were acquired between 818 and 1412 generations ago (Supplementary Fig. 29), corresponding to more than 117 passages, which is long before the extraction of the undefined starter culture in the 70s (∼27 passages). This illustrates longlasting interactions between these two phages and their bacterial hosts.

Bacterial metagenomes are biased against phage particles due to the DNA extraction method. Therefore, we cannot exclude that additional lytic phages were present in the analyzed cheese starter cultures. In line with this, a large fraction of the identified spacers matched sequences present in the viral RefSeq database (46%) including phages from all four major *Streptococcus* phage types (Fig. 6E/F). These spacers were distributed across the CRISPR arrays suggesting that the bacteria in the starter cultures have been experiencing continuous and ongoing phage pressure (Supplementary Fig. 28).

Surprisingly, we did not detect a clear negative correlation between spacer and protospacer abundance (Fig. 6G). Under the assumption that spacers represent resistant bacteria and protospacers active phages, this result does not support that bacteria-phage interactions in the starter cultures are governed by kill-the-winner dynamics^39^. This is in contrast to the previous finding in mesophilic starter cultures^11^. The exceedingly high numbers of phage copies suggest an ongoing proliferation of the phage most likely from a temperate source. Given the high abundance of the two phages in the starter cultures, despite the prevalence of matching CRISPR spacers, we assume that the phages can evade the CRISPR defense system^27,40,41^. We know that the type III CRISPR-Cas system can tolerate silent prophages^41^. Moreover, both phage genomes encode anti-CRISPR (Acr) proteins AcrIIA3^42^ and AcrIIA6^43^ known to inhibit type II CRISPR-Cas systems (Fig. 6A). Also the annotated tracrRNA of the type II-A CRISPR-Cas system has recently been shown to cause mitigation of autoimmunity but at the cost of reduced bacteriophage immunity^44^. Alternatively, the two phages may follow a carrier state or chronic infection lifestyle, in both cases the CRISPR defensive system would not be effective, as the phage particles persist either absorbed to the bacterial cell envelope (carrier state)^45^ or within the bacterial cytoplasm (chronic infection)^46,47^.

In summary, our results show that two phages are consistently present in the analyzed starter culture and outnumber bacterial cell counts indicating active phage replication without evident impact on the stability or phenotypic properties of the community.

## Discussion

Our results show that Swiss hard cheese starter cultures are dominated by a simple and highly stable microbial community. The only two detectable bacterial species were *Streptococcus thermophilus* and *Lactobacillus delbrueckii subsp. lactis*. Genomics, metabolomics and growth analysis suggest that they engage in similar metabolic interactions as found in yogurt fermentation^20^. However, positive growth effects were only observed for S. *thermophilus* on *L. delbrueckii*, but not vice versa. A more detailed analysis of their growth in co-culture will be needed to determine to what extent they cooperate, i.e. have reciprocal benefits on each other.

Strain-level diversity may contribute important functional properties to undefined cheese starter cultures, such as flavor volatile diversity, or community resilience to phage invasion or environmental disturbances^3^. We found relatively little strain-level diversity in the analyzed cheese starter cultures when compared to other microbiomes^23,48,49^. This is probably because cheese starter cultures are closed ecosystems with little opportunities for microbial migration once isolated and reproduced in a laboratory setting, as well as due to the strong population bottlenecks imposed during culture propagation. For *L. delbrueckii*, almost no genetic diversity was detected, while for *S. thermophilus* several genetically distinct variants co-existed across the analyzed samples. Cultured isolates of these strains exhibited highly similar acidification rates, growth characteristics, and volatile profiles suggesting that they are functionally redundant in phenotypic traits relevant for the cheese making process. Redundancy can promote the stability of microbial communities against perturbations^50^ and hence could explain why undefined starter cultures remain phenotypically stable.

However, why strain-level diversity is maintained in cheese starter cultures seems unclear, in particular when considering the aforementioned population bottlenecks and the fact that closely related bacteria compete for the same nutrients. Our genomic analysis revealed that in both species most of the genomic variation between strains is found in CRISPR spacers. Diverse spacers were found across the isolated strains, matching phages present in the starter cultures as well as diverse phages deposited in the reference database. Moreover, new spacers were acquired during the course of our propagation experiment. Based on these results, we conclude that undefined starter cultures must be continuously exposed to phages and suggest that strain diversity is shaped by bacteria-phage interactions.

While our metagenomic preparation was not targeted towards the sequencing of viral particles, we identified two highly abundant *Streptococcus* phages in our dataset. These phages showed almost no genetic variation across the analyzed samples (spanning hundreds of bacterial generations) suggesting that bacteria and phages do not engage in an arms-race^51,52^. Moreover, based on the correlations of protospacer-spacer abundances, we see no evidence for kill-the-winner dynamics between the two phages and *S. thermophilus*^11,53^, and hence conclude that the two phages unlikely play a role in maintaining the bacterial diversity via negative frequency dependent selection. Instead, we observed that both phages are integrated into the host genome in a subset of the cells of the different lineages, speaking in favor of a piggy-back-the-winner strategy.

Both phages seem to be maintained at high abundance in the starter cultures, without any obvious impact on bacterial growth or the cheese making process and despite the presence of a large number of matching CRISPR spacers in the bacterial population. Together, these findings suggest that the two phages have longstanding associations with their bacterial hosts and exhibit a lifestyle that may be different from other temperate phages. While phages are generally seen as a potential risk for starter cultures, it is conceivable that domesticated phages such as those found here, could also have beneficial functions. They could provide protection against lytic phages via superinfection exclusion^54^, or protect against susceptible bacterial invaders. By entering the lytic cycle at environmentally less favourable time points for the bacteria^55^, such as suboptimal pH^56^ or low nutrient ^57^ conditions, the phages may also play a direct role in cheese making as bacterial lysis is believed to enhance the flavour profile of cheese due to the liberation of intracellular proteases^58,59^.

Our work shows that domesticated microbial communities provide fascinating systems to learn about eco-evolutionary dynamics of phage-bacteria interactions. Future studies on the ecology of these phages and their role in the cheese making process could help to design artificial microbial communities in food biotechnology and beyond.

## Methods

### Strain isolation and bacterial counts

Cheese starter cultures (Lot 22.01.2019) were plated on SPY9.3 for *S. thermophilus* ^60^ and on MR11 (MRS adjusted to pH 5.4 according to ISO7889) for *L. delbrueckii*. Ninety-six colonies per species and per culture were randomly picked and inoculated into liquid media and incubated for 24 h at 37 °C. For genotyping, DNA from 100 µL of culture was extracted using the EtNaL of culture was extracted using the EtNa DNA isolation method^61^ and a minisatellite PCR was prepared as described in the supplementary methods. Isolates were stored at −40 °C in milk. Colony forming units (CFU/ml) were determined by serial dilution and plating with an Eddy jet spiral plater with counting by a SphereFlash Automatic Colony Counter (both from IUL, Barcelona, Spain) on SPY9.3 and MRS agar.

### Propagation experiment

The samples were propagated to simulate the production of cheese starter cultures. We conducted two passages per week. On Monday, 100 ul of freeze dried sample was inoculated into 10ml autoclaved biomilk media (BM) and incubated for 18 h at 37 °C. For the second passage on Tuesdays, the pre-culture was inoculated into 10ml autoclaved BM and incubated for another 18 h at 37 °C. For the final step, 100 ul of the incubated samples were transferred into a freeze dry ampulle and stored at −30 °C for at least 1 h. Thereafter, the samples were freeze dried for 7 h until dry. This procedure was repeated six times resulting in 12 propagations for five independent replicates. The number of microbial generations per incubation step were based on CFUs determined at the start and the end of the incubation, ignoring any potential cell lysis. The following formula was used to calculate the number of generations and survival rate in the freeze dry process:

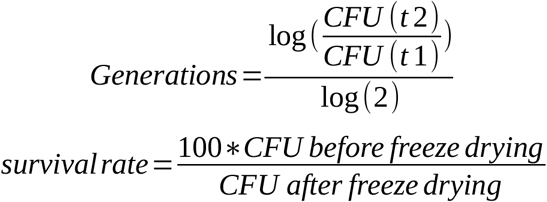

### pH measurements

For the pH measurements we used the hydroplate system (PreSens, Germany). The pH was normalized with pH standards pH 4 and 7. The measurements were done in four replicates for 30 h at 37 °C.

### Historical starter culture data

The acidification and lactate values were gathered from the Agroscope (Liebefeld, Bern) starter culture production archive. Lactate values were irregularly (approximately every 1-2 months) measured by an enzymatic essay previously described^62^.

### Genome and metagenome sequencing

Eleven samples of the cheese starter culture RMK202, including historic freeze dried ampules, present working stocks, cheese starter cultures and propagation experiment samples were prepared for shotgun metagenome sequencing. We also prepared 22 and 17 isolates of *S. thermophilus* and *L. delbrueckii* for genome sequencing, respectively. The DNA was isolated as previously explained^63^, and Nextera flex Illumina libraries prepared, (Illumina, San Diego, CA, USA) and subjected to Illumina HiSeq4000 150PE sequencing at the Genomic Technologies Facility in Lausanne, Switzerland. and with a rapid barcoding kit on a minION at the IFIK in Bern, Switzerland.

### Raw read analysis and reference genome analysis

The raw reads were trimmed with trimgalore^64^). The reads were mapped with bwa mem^65^. For the SNV-calling, freebayes-parallel was used^66^. The genomes and metagenomes were assembled with SPAdes^67^ and Flye^68^. The Flye assemblies were polished with four rounds of Racon^69^ polishing and four rounds of freebayes polishing^66^. The ANI values were calculated with fastANI^70^. The circular plots were created with circos^71^. Additionally, Panaroo^72^ and SNPeffect^73^ were used to identify core synonymous/non-synonymous mutations. The completeness of the metagenome assemblies was checked with Quast^74^, mOTU2^75^ and Busco^76^. pN/pS ratio was calculated with POGENOM^77^. The assemblies were submitted to NCBI and annotated with PGAP^78^. The core genome nucleotide phylogenies were constructed as previously described^23^

### SNV calling and strain quantification

For the single nucleotide analysis (SNV) freebayes-parallel was used with the *pooled-continuous* option for metagenomes and the following parameters: “*-C 5 --pooled-continuous -- min-alternate-fraction 0*.*05 --min-coverage 10”*. Additionally, to filter the SNVs we used vcftools^79^ with the following parameters: “*--minQ 30 --remove-indels --recode --recode-INFO-all*”. Further, only non-synonymous and SNVs in single-copy-genes were considered. The strains were quantified by using the mean alternative allele frequency of the SNVs detected solely in one lineage (For details see script).

### GC-MS

The metabolic analysis of single *S. thermophilus* and *L. delbrueckii* isolates, pairwise co-cultures, multi-strain co-cultures, and the original starter culture was done after 24h incubation in BM at 37°C. The samples for DHS-VTT-GC-MS analyses were prepared and extracted as previously described by dynamic headspace vacuum transfer in trap extraction (DHS-VTT)^80^ and analyzed by gas-chromatography mass spectrometry (GC-MS). For details see supplementary methods.

### Phage analysis

The non-bacterial reads of all metagenomes were assembled with metaSPAdes, merged and demultiplexed with cd-hit^81^. The contig coverage was estimated with bedtools^82^ and the contigs were manually curated with bandage^83^ to obtain full viral genomes (Supplementary Fig. 20). The viral genomes were annotated with Virsorter^84^, Phaster^85^ and blastn^86^. The phage network was assessed with Vcontact2^87^. We included all previously described *Streptococci* phages^88^. The location and fraction of integrated phages was assessed by identifying metagenomic paired end reads that map to both phages and bacterial genomes by filtering based on the primary and mate pair mapping location (see script). The fraction was calculated by taking the coverage of these reads in comparison to the mean coverage of the entire contig.

### CRISPR analysis

The CRISPR arrays were annotated with PilerCR^89^ and the cas genes with CRISPRcasfinder^90^. The annotation plot was done with genoplotR^91^ and the CRISPR repeat identity with the Weblogo online server^92^. Further the metagenomic spacers were extracted with a bash script (see supplementary script) and the raw spacers were demultiplexed and rarefied with DADA2^93^. CRISPR spacer turnover rate calculations were calculated by dividing the number of new CRISPR spacers in propagation experiment samples by the number of generations in that sample (∼123 generations, see Fig. 2C). Shared CRISPR spacers are calculated by clustering all extracted CRISPR spacers at 90% Identity with cd-hit-est. Metagenomic CRISPR spacers are extracted from the metagenome by using cutadapt^94^ to fish out repeat-flanking CRISPR spacers. The protospacer/spacer mapping was done by creating a complete spacer fasta database of repeat-spacer-repeat sequences. The metagenomic raw reads were mapped to this database with bwa mem^95^ and the protospacer abundance was quantified by taking the reads that mapped only to the spacer sequence. The spacer abundance was quantified by taking the reads that mapped to the complete repeat-spacer-repeat sequence

### Analysis and code

All analyses, statistics and plotting was done in R (R Core Team, 2020) and ggplot2^96^. The complete code is available at: (https://github.com/Freevini/RMK202_analysis). The two MAGs and all *S. thermophilus* and *L. delbrueckii* genomes were deposited under the NCBI Bioprojects PRJNA589532, PRJNA589608 and PRJNA659704.All data used in the analysis is available on zenodo (10.5072/zenodo.715348).

## Acknowledgments

We thank Alexandra Roetschi, Lauriane Braillard, Monika Haueter and Noam Shani for the help with the genotyping and strain collection. Anne Guisolan, Vincent Beuret and Fabio Grasso for their help in the lab. Aline Cuénod, Germán Bonilla-Rosso and Florent Mazel for critical reading of the manuscript and discussions and Joel Neugebauer for the pictogram. We also want to thank Julien Marquis and Johann Weber from the Genomic Technologies Facility in Lausanne. The project was funded by Agroscope (Liebefeld, Switzerland).

## Author contribution

VS designed and executed the project and wrote the manuscript. HB, RS and H-PB designed the project. YM and PF did the metabolomics analysis. UvA designed and advised the project. PE designed the project and wrote the manuscript. All authors gave feedback on the manuscript.

## References

1. Thompson, L. R. et al.. A communal catalogue reveals Earth’s multiscale microbial diversity. Nature 551, 457–463 (2017).

2. Nemergut, D. R. et al.. Patterns and Processes of Microbial Community Assembly. Microbiol. Mol. Biol. Rev. 77, 342–356 (2013).

3. Cogan, T. M. & Hill, C. Cheese Starter Cultures. in Cheese: Chemistry, Physics and Microbiology 193–255 (Springer, Boston, MA, 1993).

4. Powell, I. B., Broome, M. C. & Limsowtin, G. K. Y. Starter Cultures: General Aspects. Reference Module in Food Science (2016) doi:10.1016/b978-0-08-100596-5.00689-2.

5. Powell, I. B., Broome, M. C. & Limsowtin, G. K. Y. Cheese Starter Cultures: General Aspects. Encyclopedia of Dairy Sciences 261–268 (2002) doi:10.1016/b0-12-227235-8/00061-4.

6. Smid, E. J. & Lacroix, C. Microbe–microbe interactions in mixed culture food fermentations. Current Opinion in Biotechnology vol. 24 148–154 (2013).

7. Smid, E. J. et al.. Functional implications of the microbial community structure of undefined mesophilic starter cultures. Microb. Cell Fact. 13 Suppl 1, S2 (2014).

8. McCann, K. S. The diversity–stability debate. Nature 405, 228–233 (2000).

9. Lavelle, K. et al. A Decade of Streptococcus thermophilus Phage Evolution in an Irish Dairy Plant. Appl. Environ. Microbiol. 84, (2018).

10. Garneau, J. E. & Moineau, S. Bacteriophages of lactic acid bacteria and their impact on milk fermentations. Microb. Cell Fact. 10 Suppl 1, S20 (2011).

11. Erkus, O. et al. Multifactorial diversity sustains microbial community stability. ISME J. 7, 2126–2136 (2013).

12. Parente, E. et al. Microbial community dynamics in thermophilic undefined milk starter cultures. Int. J. Food Microbiol. 217, 59–67 (2016).

13. De Filippis, F., La Storia, A., Stellato, G., Gatti, M. & Ercolini, D. A selected core microbiome drives the early stages of three popular italian cheese manufactures. PLoS One 9, e89680 (2014).

14. Moser, A. et al. Amplicon Sequencing of the slpH Locus Permits Culture-Independent Strain Typing of Lactobacillus helveticus in Dairy Products. Frontiers in Microbiology vol. 8 (2017).

15. Goyal, A., Bittleston, L. S., Leventhal, G. E., Lu, L. & Cordero, O. X. Interactions between strains govern the eco-evolutionary dynamics of microbial communities. BioRxiv doi:10.1101/2021.01.04.425224.

16. Somerville, V. et al. Long-read based de novo assembly of low-complexity metagenome samples results in finished genomes and reveals insights into strain diversity and an active phage system. BMC Microbiol. 19, 143 (2019).

17. Duar, R. M. et al. Lifestyles in transition: evolution and natural history of the genus Lactobacillus. FEMS Microbiol. Rev. 41, S27–S48 (2017).

18. Bachmann, H., Starrenburg, M. J. C., Molenaar, D., Kleerebezem, M. & van Hylckama Vlieg, J. E. T. Microbial domestication signatures of Lactococcus lactis can be reproduced by experimental evolution. Genome Res. 22, 115–124 (2012).

19. Sieuwerts, S., de Bok, F. A. M., Hugenholtz, J. & van Hylckama Vlieg, J. E. T. Unraveling microbial interactions in food fermentations: from classical to genomics approaches. Appl. Environ. Microbiol. 74, 4997–5007 (2008).

20. Sieuwerts, S. et al. Mixed-culture transcriptome analysis reveals the molecular basis of mixed-culture growth in Streptococcus thermophilus and Lactobacillus bulgaricus. Appl. Environ. Microbiol. 76, 7775–7784 (2010).

21. Smit, G., Smit, B. A. & Engels, W. J. M. Flavour formation by lactic acid bacteria and biochemical flavour profiling of cheese products. FEMS Microbiology Reviews vol. 29 591– 610 (2005).

22. Schloissnig, S. et al. Genomic variation landscape of the human gut microbiome. Nature 493, 45–50 (2013).

23. Ellegaard, K. M. & Engel, P. Genomic diversity landscape of the honey bee gut microbiota. Nat. Commun. 10, 446 (2019).

24. Quince, C. et al. DESMAN: a new tool for de novo extraction of strains from metagenomes. Genome Biology vol. 18 (2017).

25. Schmid, M. et al. Comparative Genomics of Completely Sequenced Lactobacillus helveticus Genomes Provides Insights into Strain-Specific Genes and Resolves Metagenomics Data Down to the Strain Level. Front. Microbiol. 9, 63 (2018).

26. McDonnell, B. et al. A cell wall-associated polysaccharide is required for bacteriophage associated polysaccharide is required for bacteriophage adsorption to the Streptococcus thermophilus cell surface. Molecular Microbiology vol. 114 31–45 (2020).

27. Szymczak, P. et al. Cell Wall Glycans Mediate Recognition of the Dairy Bacterium Streptococcus thermophilus by Bacteriophages. Appl. Environ. Microbiol. 84, (2018).

28. Chesson, P. Mechanisms of Maintenance of Species Diversity. Annual Review of Ecology and Systematics (2000) doi:10.1146/annurev.ecolsys.31.1.343.

29. Martínez Arbas, S. et al. Roles of bacteriophages, plasmids and CRISPR immunity in microbial community dynamics revealed using time-series integrated meta-omics. Nat Microbiol 6, 123–135 (2021).

30. Makarova, K. S., Wolf, Y. I. & Koonin, E. V. The basic building blocks and evolution of CRISPR–Cas systems. Biochemical Society Transactions vol. 41 1392–1400 (2013).

31. Hynes, A. P., Villion, M. & Moineau, S. Adaptation in bacterial CRISPR-Cas immunity can be driven by defective phages. Nat. Commun. 5, 4399 (2014).

32. Kanmani, P. et al. Genomic Characterization of Lactobacillus delbrueckii TUA4408L and Evaluation of the Antiviral Activities of its Extracellular Polysaccharides in Porcine Intestinal Epithelial Cells. Frontiers in Immunology vol. 9 (2018).

33. Shmakov, S. A. et al. The CRISPR Spacer Space Is Dominated by Sequences from Species-Specific Mobilomes. MBio 8, (2017).

34. Paez-Espino, D. et al. CRISPR immunity drives rapid phage genome evolution in Streptococcus thermophilus. MBio 6, (2015).

35. Villanueva, V. M., Oldfield, L. M. & Hatfull, G. F. An Unusual Phage Repressor Encoded by Mycobacteriophage BPs. PLoS One 10, e0137187 (2015).

36. Ruiz-Cruz, S. et al. Lysogenization of a Lactococcal Host with Three Distinct Temperate Phages Provides Homologous and Heterologous Phage Resistance. Microorganisms 8, (2020).

37. Szymczak, P. et al. Novel Variants of Streptococcus thermophilus Bacteriophages Are Indicative of Genetic Recombination among Phages from Different Bacterial Species. Appl. Environ. Microbiol. 83, (2017).

38. Staals, R. H. J. et al. Interference-driven spacer acquisition is dominant over naive and primed adaptation in a native CRISPR–Cas system. Nature Communications vol. 7 (2016).

39. Weitz, J. S., Beckett, S. J., Brum, J. R., Cael, B. B. & Dushoff, J. Lysis, lysogeny and virus– microbe ratios. Nature vol. 549 E1–E3 (2017).

40. Common, J., Morley, D., Westra, E. R. & van Houte, S. CRISPR-Cas immunity leads to a coevolutionary arms race between Streptococcus thermophilus and lytic phage. Philos. Trans. R. Soc. Lond. B Biol. Sci. 374, 20180098 (2019).

41. Li, Y. & Bondy-Denomy, J. Anti-CRISPRs go viral: the infection biology of CRISPR-Cas inhibitors. Cell Host Microbe (2021) doi:10.1016/j.chom.2020.12.007.

42. Rauch, B. J. et al. Inhibition of CRISPR-Cas9 with Bacteriophage Proteins. Cell vol. 168 150–158.e10 (2017).

43. Fuchsbauer, O. et al. Cas9 Allosteric Inhibition by the Anti-CRISPR Protein AcrIIA6. Mol. Cell 76, 922–937.e7 (2019).

44. Workman, R. et al. A natural single-guide RNA repurposes Cas9 to autoregulate CRISPR-Cas expression. Cell (2021) doi:10.1016/j.cell.2020.12.017.

45. Guerin, E. & Hill, C. Shining Light on Human Gut Bacteriophages. Front. Cell. Infect. Microbiol. 10, (2020).

46. Roux, S. et al. Cryptic inoviruses revealed as pervasive in bacteria and archaea across Earth’s biomes. Nature Microbiology 4, 1895–1906 (2019).

47. Shkoporov, A. N. et al. Long-term persistence of crAss-like phage crAss001 is associated with phase variation in Bacteroides intestinalis. (2020) doi:10.1101/2020.12.02.408625.

48. Gregory, A. C. et al. Marine DNA Viral Macro-and Microdiversity from Pole to Pole. Cell 177, 1109–1123.e14 (2019).

49. Nayfach, S. et al. A genomic catalog of Earth’s microbiomes. Nat. Biotechnol. 1–11 (2020).

50. Coyte, K. Z., Schluter, J. & Foster, K. R. The ecology of the microbiome: Networks, competition, and stability. Science 350, 663–666 (2015).

51. Rodriguez-Valera, F. et al. Explaining microbial population genomics through phage predation. Nat. Rev. Microbiol. 7, 828–836 (2009).

52. Deveau, H. et al. Phage response to CRISPR-encoded resistance in Streptococcus thermophilus. J. Bacteriol. 190, 1390–1400 (2008).

53. Thingstad, T. F. Elements of a theory for the mechanisms controlling abundance, diversity, and biogeochemical role of lytic bacterial viruses in aquatic systems. Limnology and Oceanography vol. 45 1320–1328 (2000).

54. Bondy-Denomy, J. et al. Prophages mediate defense against phage infection through diverse mechanisms. ISME J. 10, 2854–2866 (2016).

55. O’Sullivan, D., Ross, R. P., Fitzgerald, G. F. & Coffey, A. Investigation of the relationship between lysogeny and lysis of Lactococcus lactis in cheese using prophage-targeted PCR. Appl. Environ. Microbiol. 66, 2192–2198 (2000).

56. Lunde, M., Aastveit, A. H., Blatny, J. M. & Nes, I. F. Effects of Diverse Environmental Conditions on fLC3 Prophage Stability in Lactococcus lactis. LC3 Prophage Stability in Lactococcus lactis. Appl. Environ. Microbiol. 71, 721–727 (2005).

57. Matos, A. P. A. de et al. Comparison of induction of B45 Helicobacter pylori prophage by acid and UV radiation. Microscopy and Microanalysis vol. 19 27–28 (2013).

58. Smid, E. J. & Kleerebezem, M. Production of Aroma Compounds in Lactic Fermentations. Annual Review of Food Science and Technology vol. 5 313–326 (2014).

59. Nugroho, A. D. W., Kleerebezem, M. & Bachmann, H. Growth, dormancy and lysis: the complex relation of starter culture physiology and cheese flavour formation. Current Opinion in Food Science (2020) doi:10.1016/j.cofs.2020.12.005.

60. Shani, N., Isolini, D., Marzohl, D. & Berthoud, H. Evaluation of a new culture medium for the enumeration and isolation of Streptococcus salivarius subsp. thermophilus from cheese. Food Microbiology vol. 95 103672 (2021).

61. Vingataramin, L. & Frost, E. H. A single protocol for extraction of gDNA from bacteria and yeast. Biotechniques 58, 120–125 (2015).

62. Borshchevskaya, L. N., Gordeeva, T. L., Kalinina, A. N. & Sineokii, S. P. Spectrophotometric determination of lactic acid. Journal of Analytical Chemistry vol. 71 755–758 (2016).

63. Moser, A., Berthoud, H., Eugster, E., Meile, L. & Irmler, S. Detection and enumeration of Lactobacillus helveticus in dairy products. International Dairy Journal vol. 68 52–59 (2017).

64. Martin, M. Cutadapt removes adapter sequences from high-throughput sequencing reads. EMBnet.journal vol. 17 10 (2011).

65. Li, H. & Durbin, R. Fast and accurate long-read alignment with Burrows-Wheeler transform. Bioinformatics 26, 589–595 (2010).

66. Garrison, E. & Marth, G. Haplotype-based variant detection from short-read sequencing. arXiv.org (2012) doi:arXiv:1207.3907.

67. Nurk, S. et al. Assembling Genomes and Mini-metagenomes from Highly Chimeric Reads. Lecture Notes in Computer Science 158–170 (2013) doi:10.1007/978-3-642-37195-0_13.

68. Kolmogorov, M., Yuan, J., Lin, Y. & Pevzner, P. A. Assembly of long, error-prone reads using repeat graphs. Nat. Biotechnol. 37, 540–546 (2019).

69. Vaser, R., Sović, I., Nagarajan, N. & Šikić, M. Fast and accurate de novo genome assembly from long uncorrected reads. Genome Research vol. 27 737–746 (2017).

70. Jain, C., Rodriguez-R, L. M., Phillippy, A. M., Konstantinidis, K. T. & Aluru, S. High throughput ANI analysis of 90K prokaryotic genomes reveals clear species boundaries. Nat. Commun. 9, 5114 (2018).

71. Krzywinski, M. et al. Circos: an information aesthetic for comparative genomics. Genome Res. 19, 1639–1645 (2009).

72. Tonkin-Hill, G. et al. Producing polished prokaryotic pangenomes with the Panaroo pipeline. Genome Biol. 21, 180 (2020).

73. De Baets, G. et al. SNPeffect 4.0: on-line prediction of molecular and structural effects of protein-coding variants. Nucleic Acids Res. 40, D935–9 (2012).

74. Mikheenko, A., Saveliev, V. & Gurevich, A. MetaQUAST: evaluation of metagenome assemblies. Bioinformatics 32, 1088–1090 (2016).

75. Milanese, A. et al. Microbial abundance, activity and population genomic profiling with mOTUs2. Nat. Commun. 10, 1014 (2019).

76. Simão, F. A., Waterhouse, R. M., Ioannidis, P., Kriventseva, E. V. & Zdobnov, E. M. BUSCO: assessing genome assembly and annotation completeness with single-copy orthologs. Bioinformatics 31, 3210–3212 (2015).

77. Sjöqvist, C., Zambrano, L. F. D., Alneberg, J. & Andersson, A. F. Revealing ecologically coherent population structure of uncultivated bacterioplankton with POGENOM. bioRxiv (2020) doi:10.1101/2020.03.25.999755.

78. Tatusova, T. et al. NCBI prokaryotic genome annotation pipeline. Nucleic Acids Research vol. 44 6614–6624 (2016).

79. Danecek, P. et al. The variant call format and VCFtools. Bioinformatics vol. 27 2156–2158 (2011).

80. Fuchsmann, P. et al. Development and performance evaluation of a novel dynamic headspace vacuum transfer ‘In Trap’ extraction method for volatile compounds and comparison with headspace solid-phase microextraction and headspace in-tube extraction. Journal of Chromatography A vol. 1601 60–70 (2019).

81. Li, W. & Godzik, A. Cd-hit: a fast program for clustering and comparing large sets of protein or nucleotide sequences. Bioinformatics vol. 22 1658–1659 (2006).

82. Quinlan, A. R. BEDTools: The Swiss-Army Tool for Genome Feature Analysis. Curr. Protoc. Bioinformatics 47, 11.12.1–34 (2014).

83. Wick, R. R., Schultz, M. B., Zobel, J. & Holt, K. E. Bandage: interactive visualization of de novo genome assemblies. Bioinformatics 31, 3350–3352 (2015).

84. Roux, S., Enault, F., Hurwitz, B. L. & Sullivan, M. B. VirSorter: mining viral signal from microbial genomic data. PeerJ 3, e985 (2015).

85. Arndt, D. et al. PHASTER: a better, faster version of the PHAST phage search tool. Nucleic Acids Res. 44, W16–21 (2016).

86. Altschul, S. F., Gish, W., Miller, W., Myers, E. W. & Lipman, D. J. Basic local alignment search tool. J. Mol. Biol. 215, 403–410 (1990).

87. Bolduc, B. et al. vConTACT: an iVirus tool to classify double-stranded DNA viruses that infect and. PeerJ 5, e3243 (2017).

88. Lavelle, K. et al. Biodiversity of Phages in Global Dairy Fermentations. Viruses 10, (2018).

89. Edgar, R. C. PILER-CR: fast and accurate identification of CRISPR repeats. BMC Bioinformatics 8, 18 (2007).

90. Couvin, D. et al. CRISPRCasFinder, an update of CRISRFinder, includes a portable version, enhanced performance and integrates search for Cas proteins. Nucleic Acids Res. 46, W246–W251 (2018).

91. Guy, L., Kultima, J. R. & Andersson, S. G. E. genoPlotR: comparative gene and genome visualization in R. Bioinformatics 26, 2334–2335 (2010).

92. Crooks, G. E. WebLogo: A Sequence Logo Generator. Genome Research vol. 14 1188– 1190 (2004).

93. Callahan, B. J. et al. DADA2: High-resolution sample inference from Illumina amplicon data. Nat. Methods 13, 581–583 (2016).

94. Martin, M. Cutadapt removes adapter sequences from high-throughput sequencing reads. EMBnet.journal 17, 10–12 (2011).

95. Li, H. & Durbin, R. Fast and accurate long-read alignment with Burrows-Wheeler transform. Bioinformatics 26, 589–595 (2010).

96. Wickham, H. ggplot2: Elegant Graphics for Data Analysis. (Springer Science & Business Media, 2009).

