## Supplemental Information for "Functional strain redundancy and persistent phage infection in Swiss hard cheese starter cultures"

#### Content overview

Supplemental methods……………………………………………………………………………….2

Supplemental figures……………………………………………………………………………..….5

### Supplemental methods

#### Genotyping

For genotyping, DNA from 100 µL of culture was extracted using the EtNa DNA isolation method (Vingataramin and Frost 2015) and diluted 10x before PCR amplification. Multiplex amplification was performed in 25 µL reactions containing 2.5 µL of GeneAmp® 10X PCR Buffer I containing 15 mM $MgCl_2$ (Thermo Fisher, Waltham, MA, United States), 0.9 mL $MgCl_2$ (50mM), 0.5 mL PCR nucleotide mix (10 mM; Promega AG, Dübendorf, Switzerland), 0.5 µL Q-Solution (Qiagen, Hombrechtikon, Switzerland), 2.5 µL primer mix (2 mM of each primer listed in SupplementTable1), 0.4 µL AmpliTaq Gold DNA polymerase (Thermo Fisher) and 2 µL of DNA.
After the initial heat activation at 95°C for 10 min followed 35 cycles at 94°C for 1 min, 58°C for 30 s, 72°C for 30 s, and a final extension at 72°C for 7 min. The amplification products were separated using the DNF-905-K0500 dsDNA 905 Reagent Kit (separation range 1-500 bp) on a Fragment Analyzer™ (Advanced Analytical Technologies, Ankeny, IA, USA) according to the manufacturer's instructions. The results were evaluated and compared with the PROSize software (V.3, Advanced Analytical Technologies).

#### GC-MS analysis

Samples for DHS-VTT-GC-MS analyses were prepared as follows: 250 mg of cheese starter culture were weighed in 20 mL headspace crimp glass vials. 25 µL of an internal standard solution composed by 0.5 ppm of paraldehyde, 0.25 ppm of tetradecane and 0.5 ppm of d4-δ-decalactone were added to the samples for analytical deviation correction.

Volatiles were extracted by dynamic headspace vacuum transfer in trap extraction (DHS-VTT) (Fuchsmann et al. 2019) and analyzed by gas-chromatography mass spectrometry (GC-MS). Volatile were adsorbed on a Tenax TA/Carbosieve III ITEX (in tube extraction) trap (BGB Analytik AG, Böckten, Switzerland) which was conditioned according to the supplier's temperature recommendations (320°C for 1 h) under a nitrogen stream of 100 mL min-1. During extraction, the syringe temperature was fixed at 100°C and the ITEX trap at 35°C. Samples were incubated for 10 min at 60°C and volatiles were extracted for 5 min at 5 mbar using a vacuum pump Buchi V-300 (Büchi, Flawil, Switzerland). After extraction, the sorbent and syringe were dried under a nitrogen stream for 5 and 20 min, respectively, at 220 mL min-1 to avoid injection of water in the column. Bound volatiles were desorbed for 2 min with a nitrogen flow of 100 mL min-1 at 240°C in a programmed temperature vaporizer (PTV) injector of type CIS4 (Gerstel AG, Sursee, Switzerland) in the vent mode at 50 mL min-1 and 0 kPa for 30 s. The injector containing a Tenax TA filled glass cooled to 10°C using liquid nitrogen trapped the compounds again. The volatiles were then released by heating the injector at a rate of 12°C sec-1 to 240°C. After injection, the trap was reconditioned according to the supplier's temperature recommendation (300°C) for 15 min under a nitrogen flow of 100 mL min-1.

The analyses were completed using an MPS2 autosampler (Gerstel AG) on an Agilent 7890B GC system coupled to an Agilent 5977B mass selective detector (MSD) (Agilent Technologies, Basel, Switzerland). Volatile compounds were separated on a OPTIMA FFAPplus fused silica capillary column (polyethylene glycol nitroterephthalate, crosslinked, 60 m × 0.25 mm × 0.5 µm film; MACHEREY-NAGEL, Düren, Germany) with helium as the carrier gas at a constant flow of 1.5 mL min-1 (25.312 cm sec-1).
The oven temperature was programmed as follows: 5 min at 40°C, then heated to 240°C at a rate of 5°C min-1 with a final hold time of 20 min to make a total run time of 65 min.
The MS settings were as follows: transfer line at 250°C, source temperature at 230°C, and the analytes monitored in SCAN mode between 30 amu and 350 amu with 4 min solvent delay. The autosampler was controlled with a Cycle Composer V.1.5.4 (CTC Analytics, Zwingen, Switzerland) and PTV injector with Maestro1 software V.1.4.8.14/3.5 (Gerstel AG).

Peak extraction and grouping was obtained using MassHunter Profinder software version 10.0 (Agilent Technologies). A batch recursive feature extraction (small molecules/peptides) was performed on all data. For the molecular feature extraction (MFE), peaks with a height lower than 2000 counts were filtered out. Alignment parameters were set with a 0.30 min retention time tolerance and a 0.40 minimal dot-product value. A post-processing filter was set for the MFE with a score of minimum 50.0 with the requirement that a compound must satisfy the condition in at least one file across all sample files. For the Find by Ion parameters, a mass tolerance was set at ± 200 mDa and a retention time window of ± 0.30 min. Agile 2 integration was chosen and the chromatograms were smoothed using the Gaussian function with the function and Gaussian width both at 7 points. Spectra extraction was set to include spectra at apex of peak.

### Supplemental figures

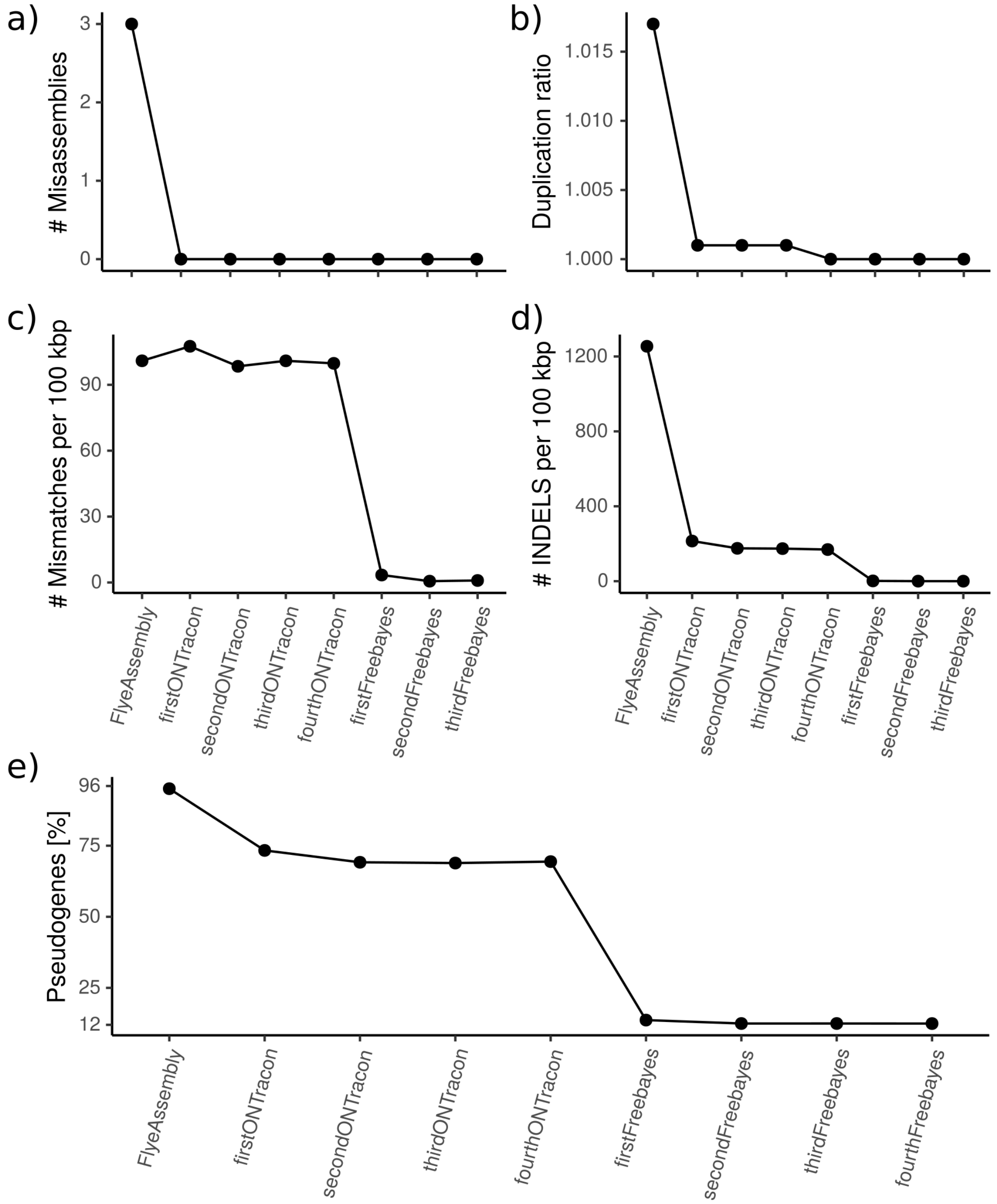


Figure 1. Polishing of the metagenome-assembled-genomes (MAGs). The quality can be illustrated by the steady decrease of a) misassemblies, b) duplication rate, c) mismatches, d) INDELS, and e) pseudogenes over the four Racon-based polishing steps and the four Freebayes-based polishing steps.

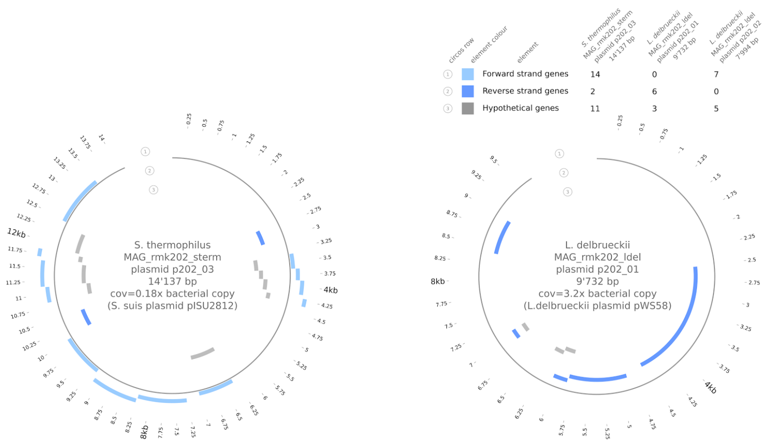


Figure 2. The metagenome assembled plasmids. The plots include gene annotations, and are labelled with plasmid name, size, coverage (relative to bacterial host), and closest blast hit.

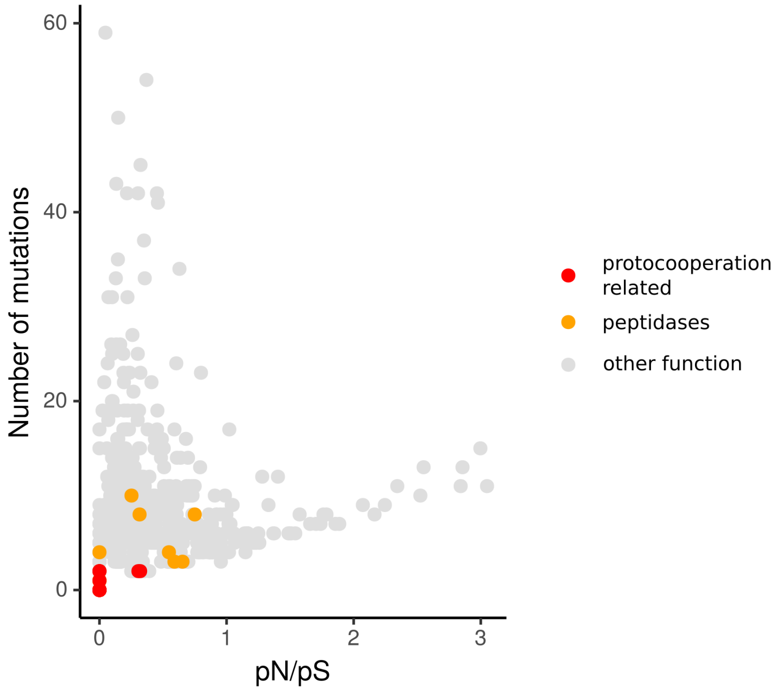


Figure 3. pN/pS ratios and number of mutations of all genes of the *L. delbrueckii* and *S. thermophilus* MAGs. The genes related to protocooperation and the peptidases are colored accordingly.

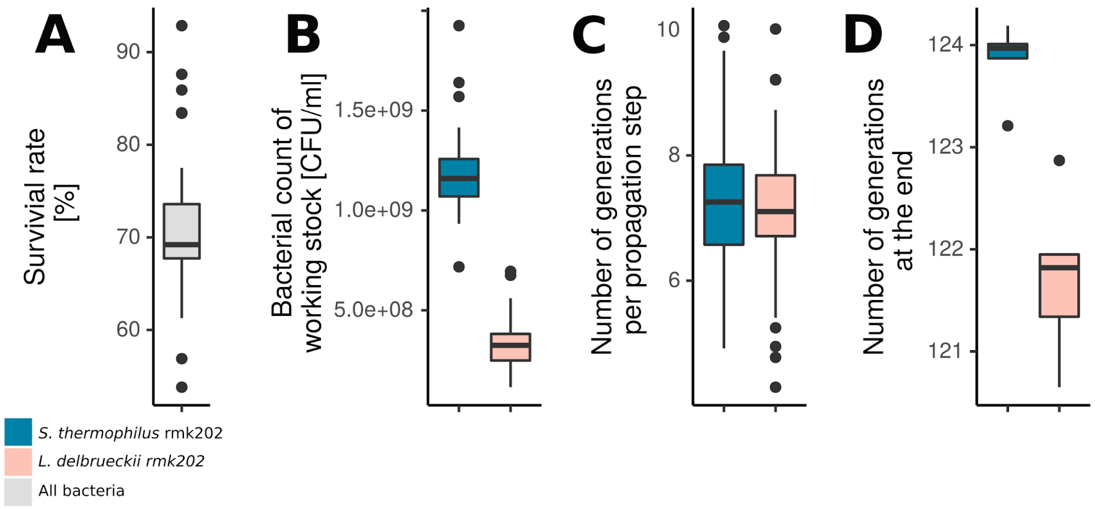


Figure 4. The analysis of the phenotypic data of the propagation experiment. A) The overall survival rate of the bacteria after freeze drying. B) The bacterial counts in CFU/ml of the working stocks for both species. C) The number of generations per passage over the entire experiment for both species. D) The final number of generations at the end of the evolution experiment per species.

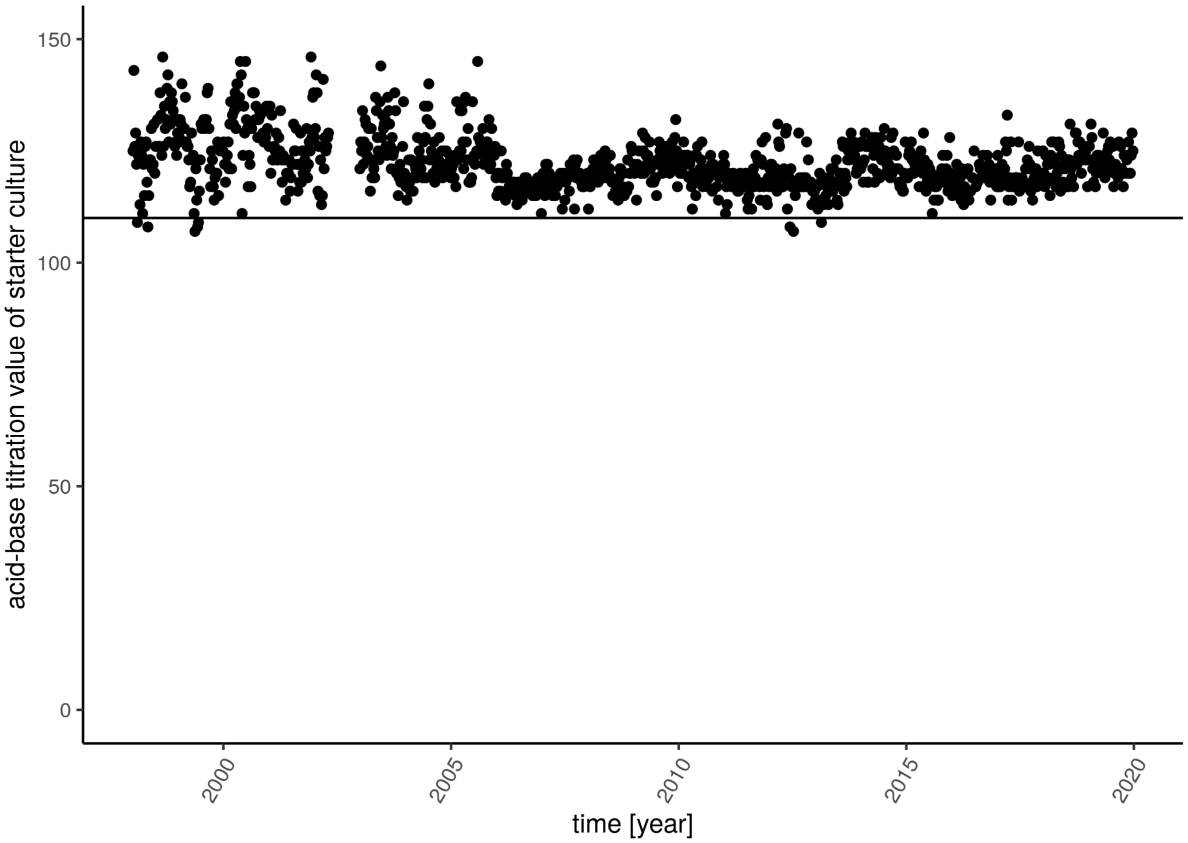


Figure 5. The acid-based titratable value of the starter culture RMK202 ranging back to 1996. The titratable acidity is measured after incubation for 18 h at 37°C in milk. The black line is the lowest minimum accepted value.

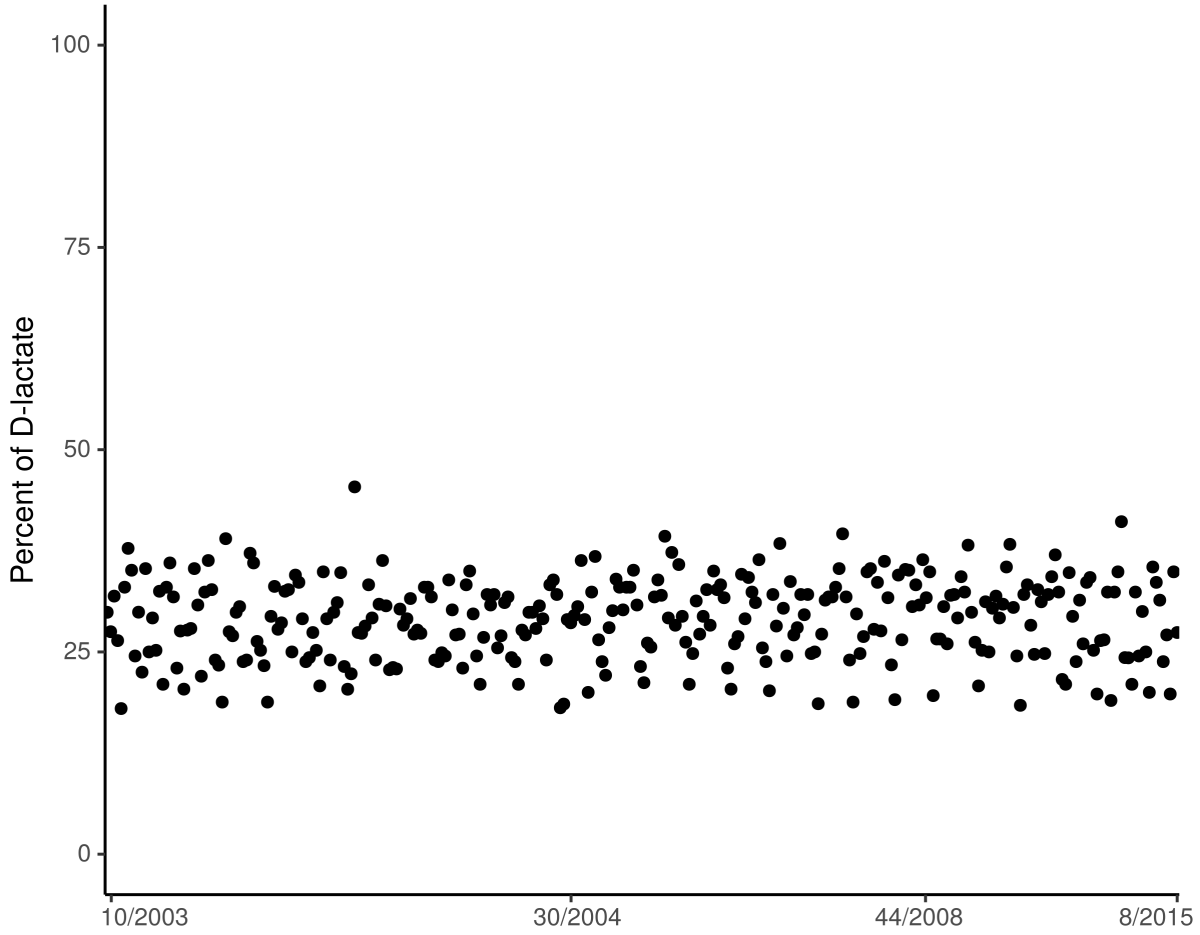


Figure 6. The percent of D-lactate to L-lactate measured after 18 h of incubation at 37°C in milk. D-lactate is produced by *L. delbrueckii* and L-lactate by *S. thermophilus*. Measurements were irregularly conducted between 2003 and 2015. Dates are illustrated on x-axis.

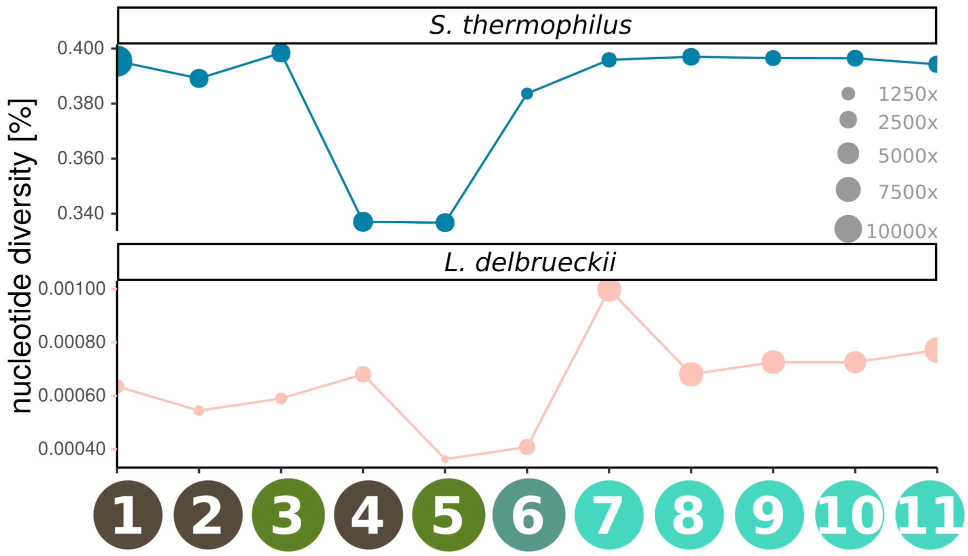


Figure 7. The fraction of variable sites (nucleotide diversity) in the housekeeping genes of the *S.thermophilus* (top) and *L. delbrueckii* (bottom) over the 11 metagenomic samples. The coverage on the individual samples is indicated with the size of the point. (The legend for the samples on the x-axis are illustrated in Fig. 2A).

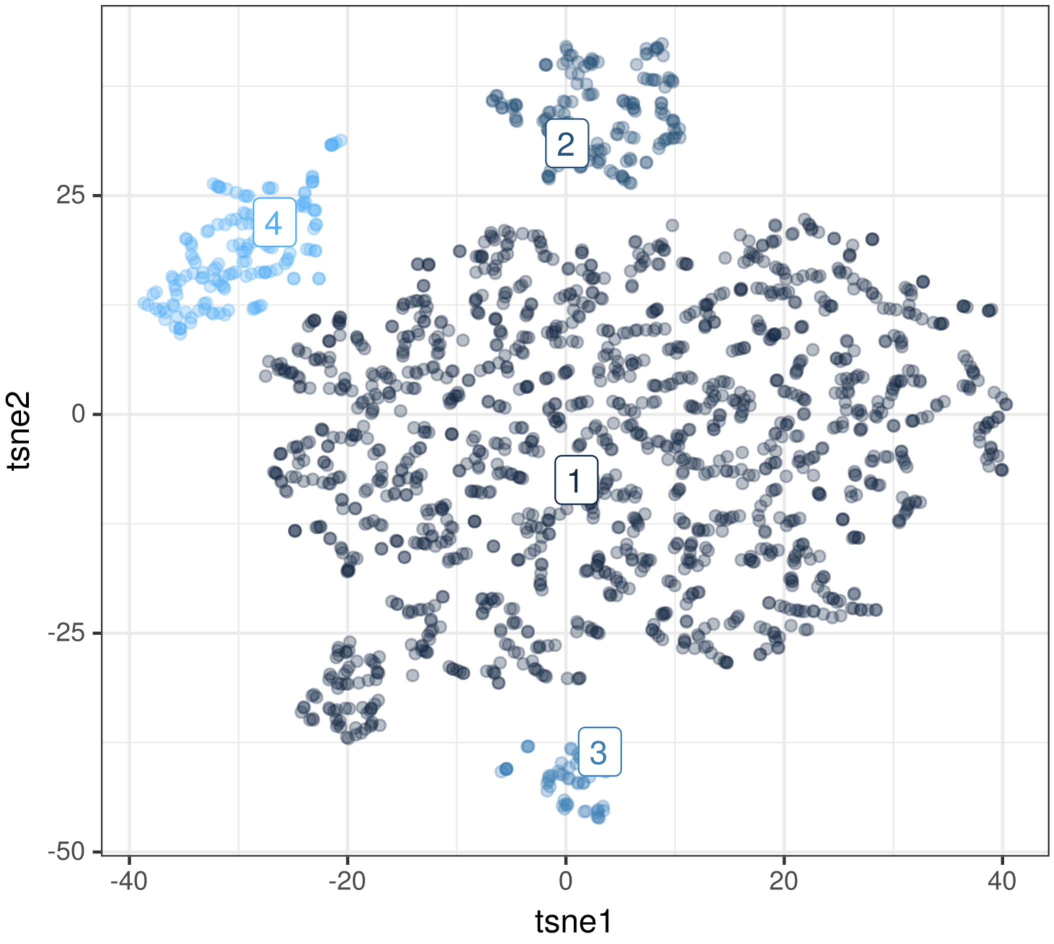


Figure 8. The Tsne clustering of all metagenomic *S. thermophilus* SNVs.

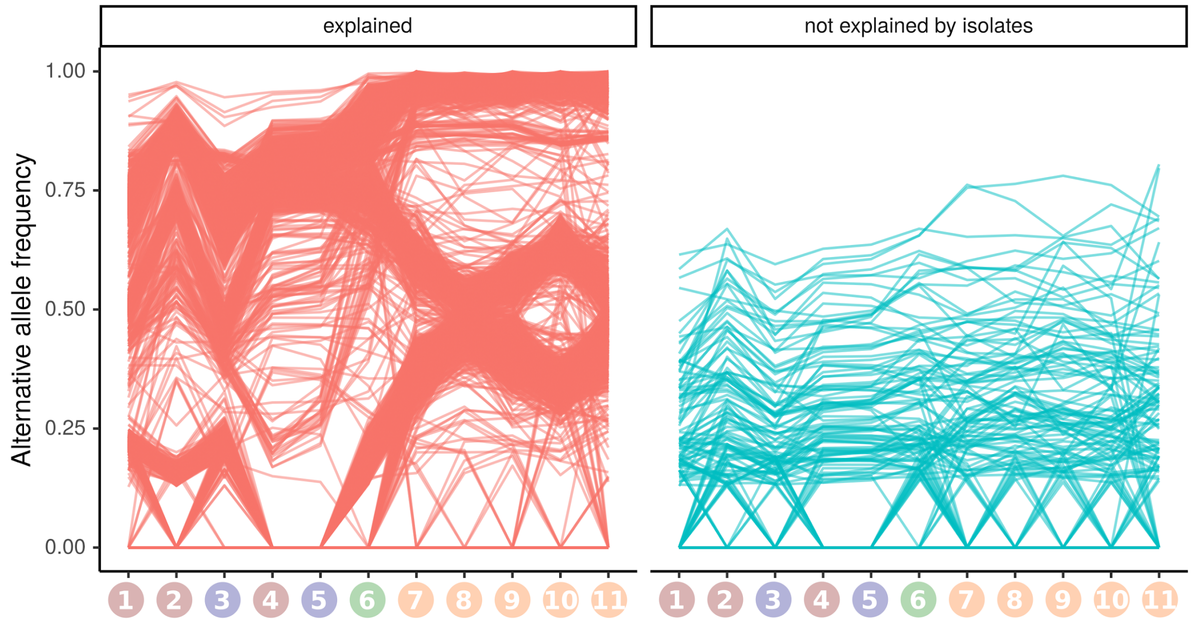


Figure 9. The alternative allele frequency of all *S. thermophilus* SNVs that are explained (97%) and not explained (3%) by the isolates. The x-axis labels correspond to the sample annotations in figure 2A.

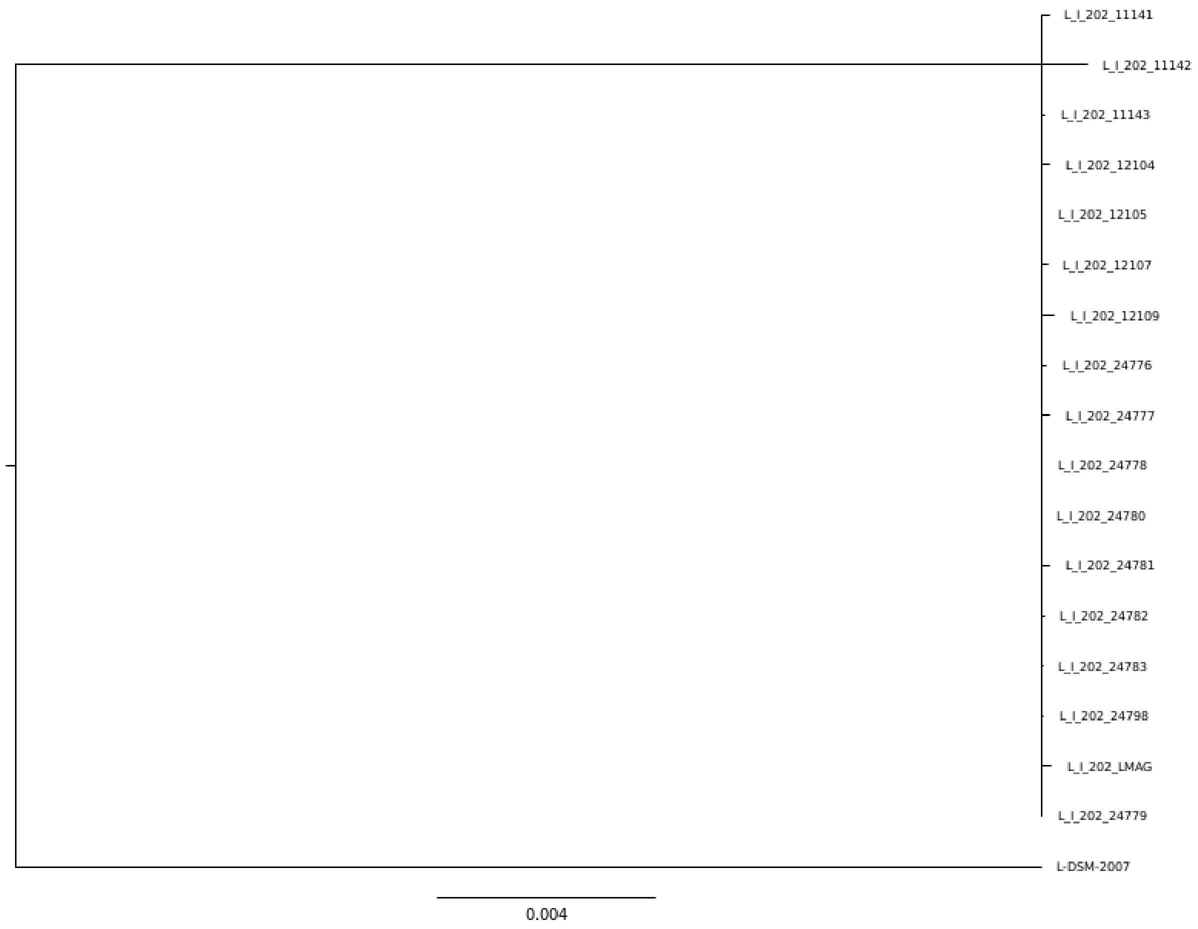


Figure 10. Phylogeny of all RMK202 *L. delbrueckii* isolates with the *L. delbrueckii subsp. lactis* type strain DSM-2007 as outgroup. The phylogeny is based on 1596 core genes.

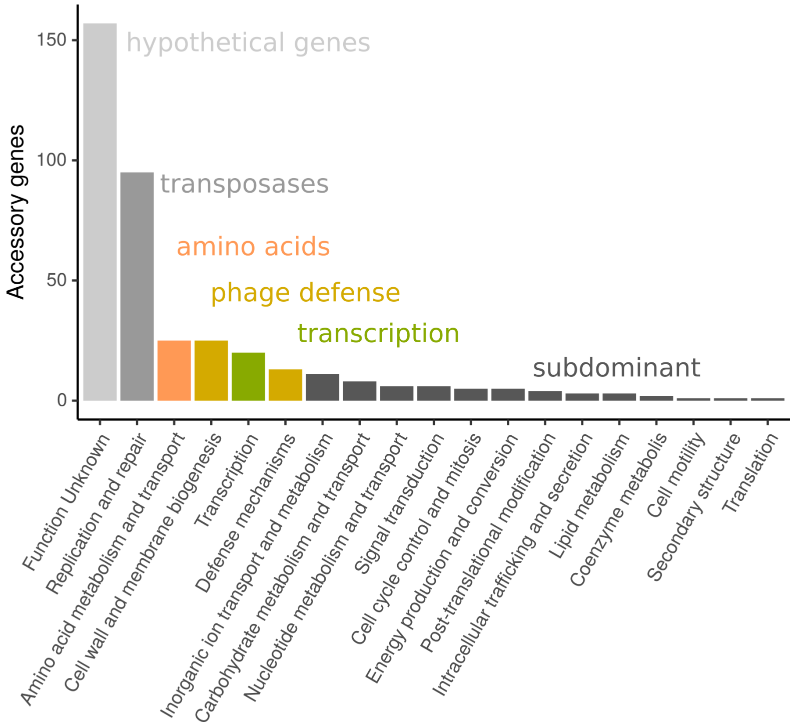


Figure 11. The accessory genes of *S. thermophilus* grouped into different COG categories.

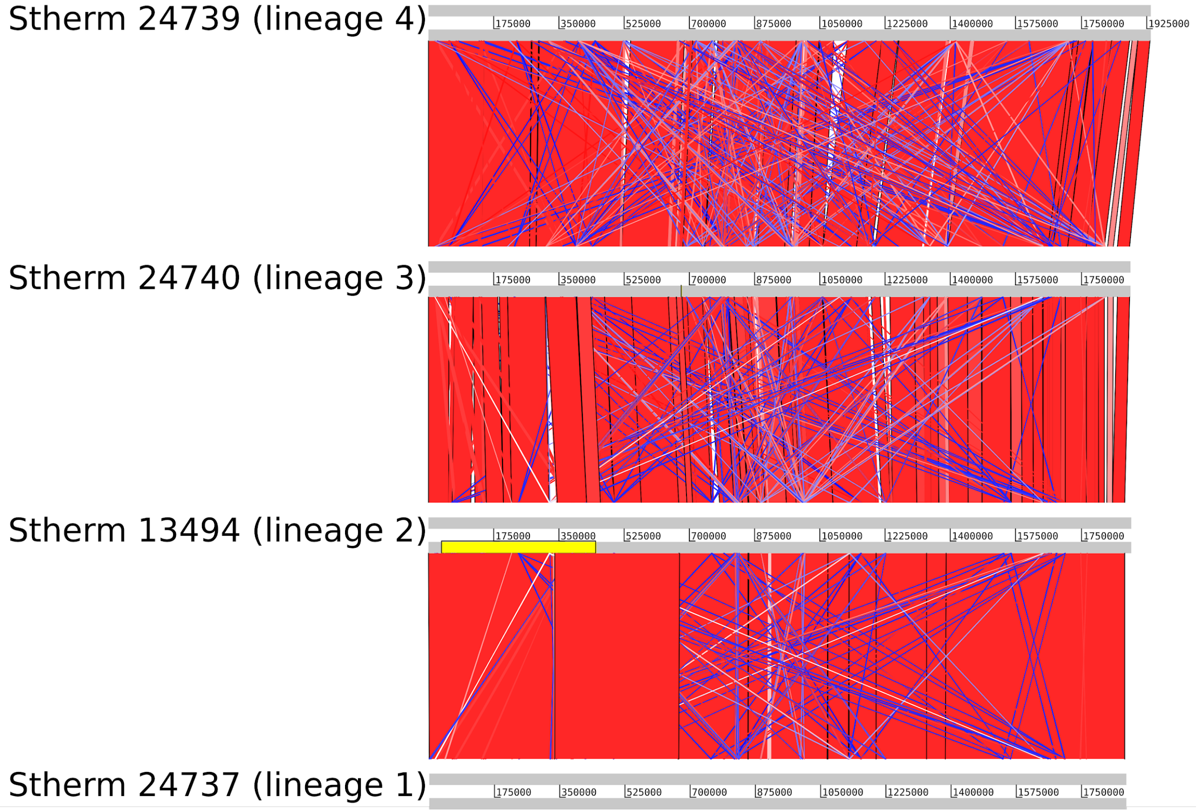


Figure 12. Artemis collinearity plot of the representative genomes from all four lineage of *S. thermophilus*. The regions are selected for min 1000bp and min 90% nucleotide identity. The collinear regions illustrated are colored according to orientation (red=same orientation, blue=reversed).

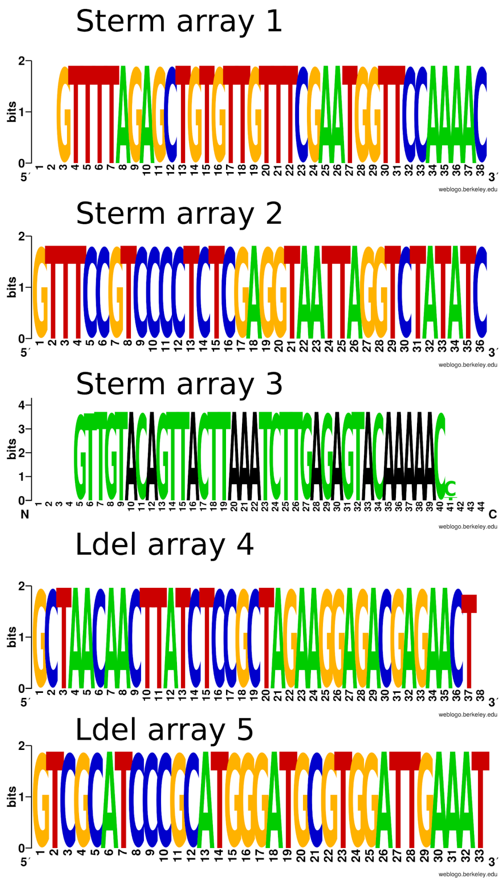


Figure 13. CRISPR repeat conservation over all assembled genomes is illustrated in the weblogo. (Sterm= *S. thermophilus* and Ldel= *L. delbrueckii)*

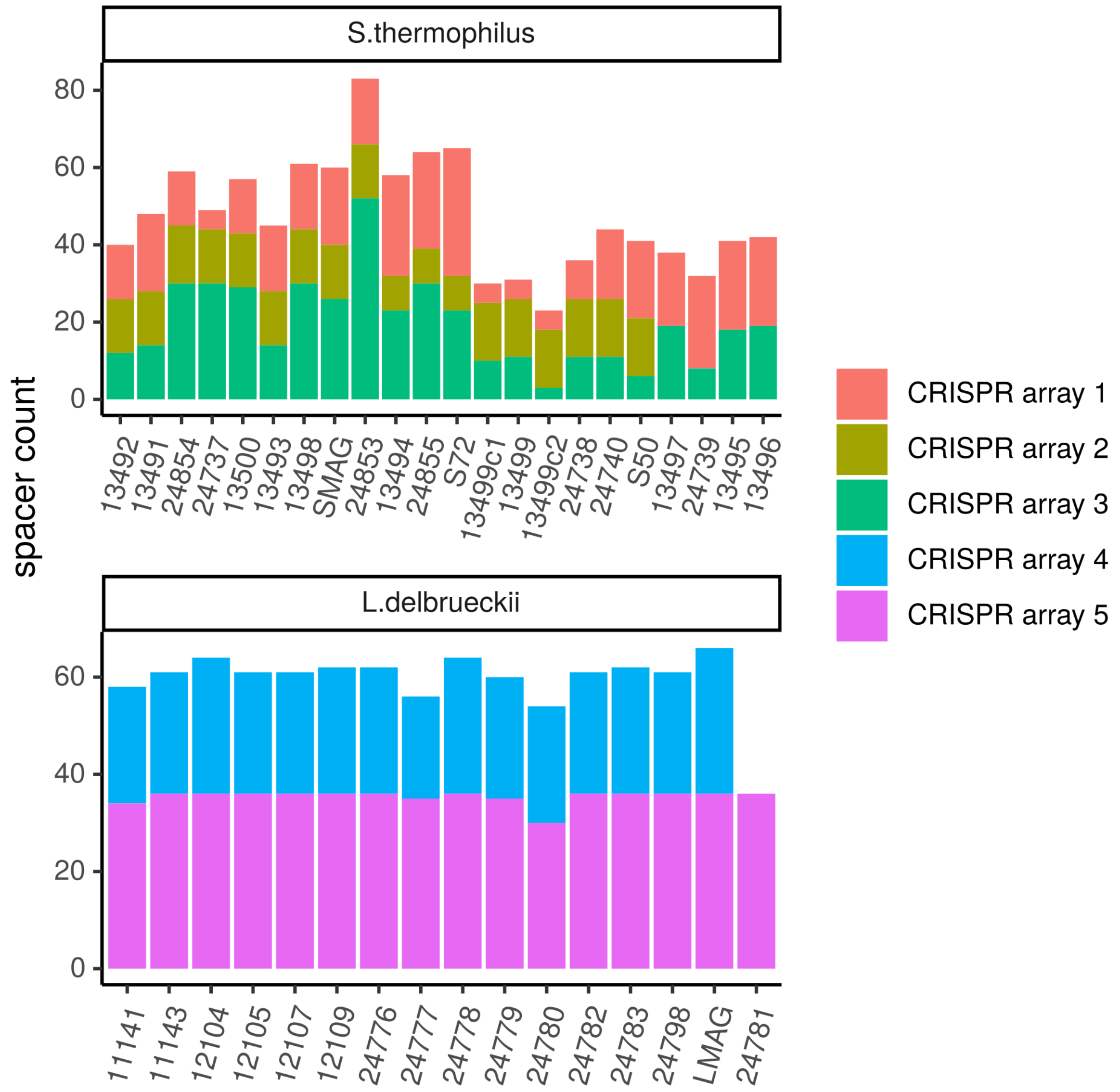


Figure 14. Number of spacers per *S. thermophilus* (top) and *L. delbrueckii* (top) strains. Colors indicate the respective CRISPR array. (*L. delbrueckii* 24781 does not contain any CRISPR array 4)

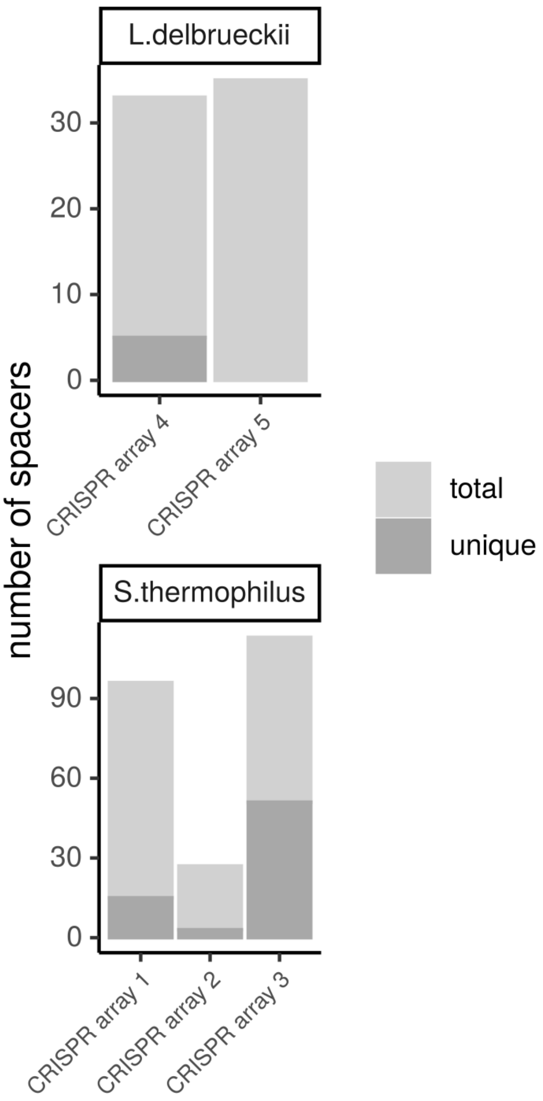


Figure 15. The overall number of spacers in the five arrays. The fraction of unique spacers are indicated in darker gray.

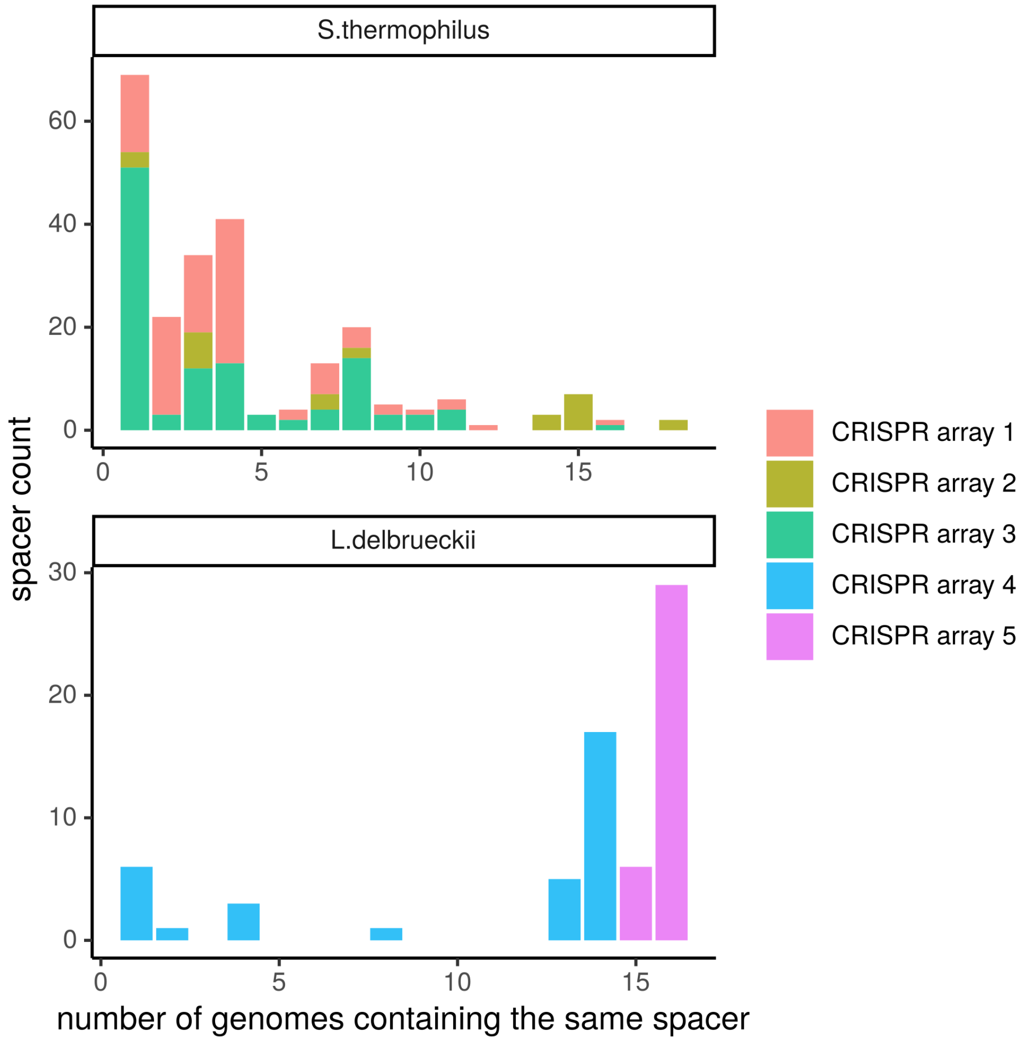


Figure 16. Number of genomes containing the same spacer. The colors indicate the CRISPR array the spacer is associated with.

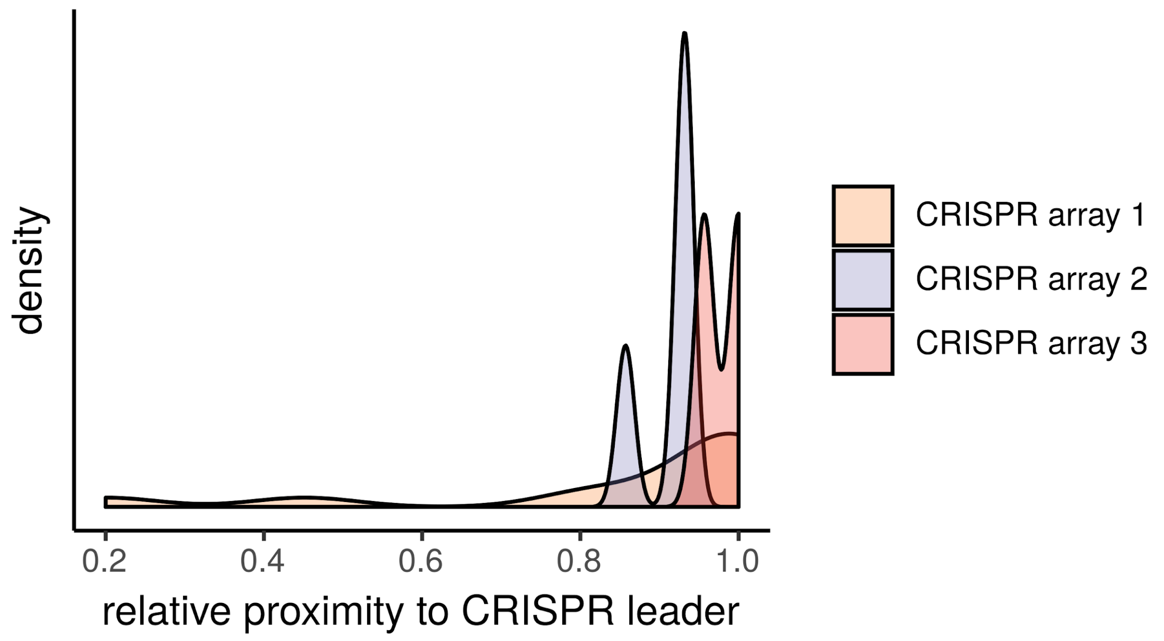


Figure 17. The relative proximity to the CRISPR leader of all unique CRISPR spacers isolated from the *S. thermophilus* genomes.

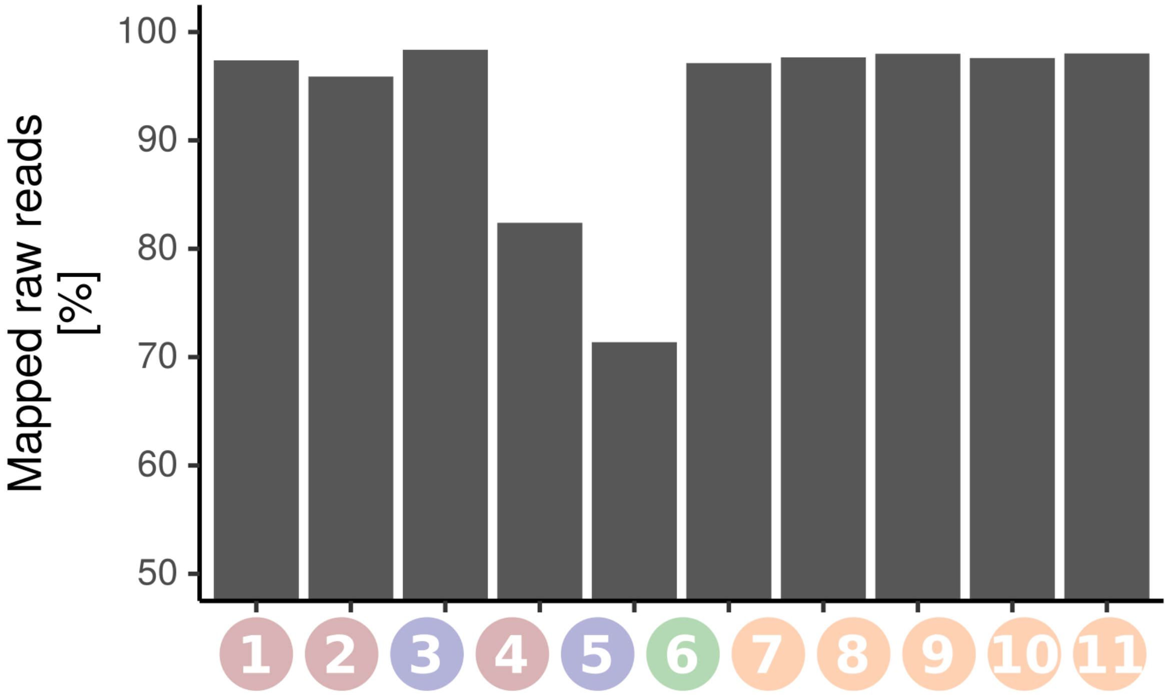


Figure 18. Percent of mapped raw reads for the different metagenomic samples against the MAGs. The sample labels on x-axis are chronologically and described in more detail in Fig. 2A.

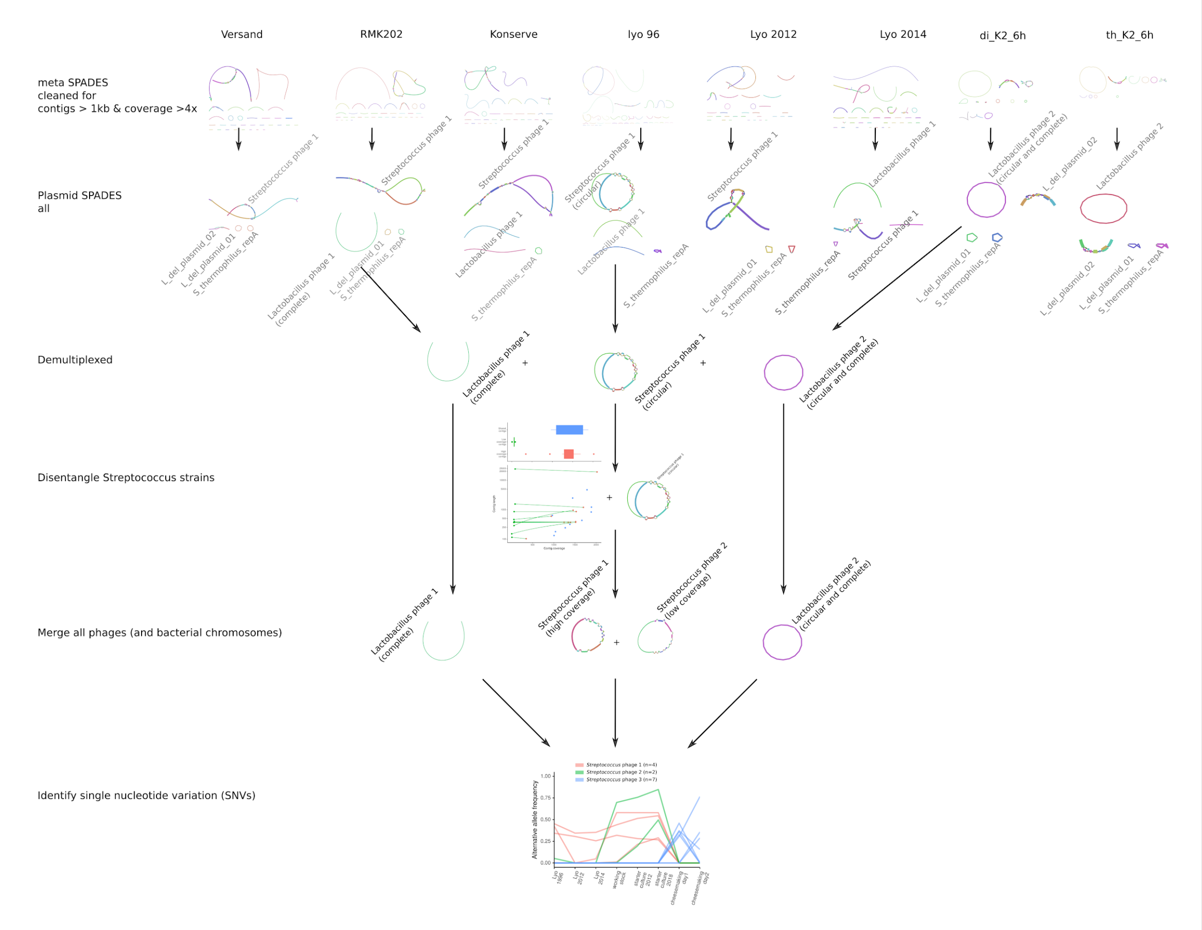


Figure 19. The curated phage assembly with all involved steps: 1) metaSPAdes assembly of unmapped reads. 2) compare with plasmidSPAdes assembly of unmapped reads 3) demultiplex all assemblies with cd-hit. 4) Disentangle *Streptococcus* phages based on contig coverage in bandage (Wick et al. 2015). 5) Merge the disentangled phages and check continuous mapping. 6) Identify single nucleotide variations (SNVs) on the curated phage genomes

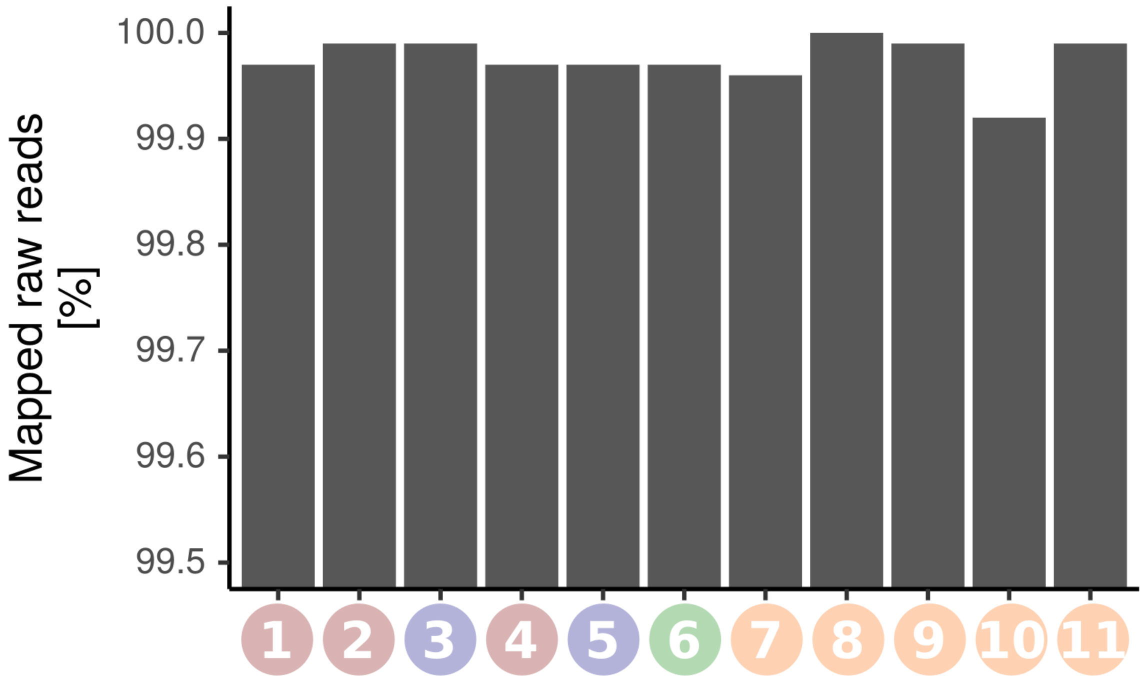


Figure 20. Percent of mapped raw reads for the different metagenomic samples (mean=99.97%, sd=0.02%). Sample labels on x-axis are chronologically and described in more detail in Fig. 2A.

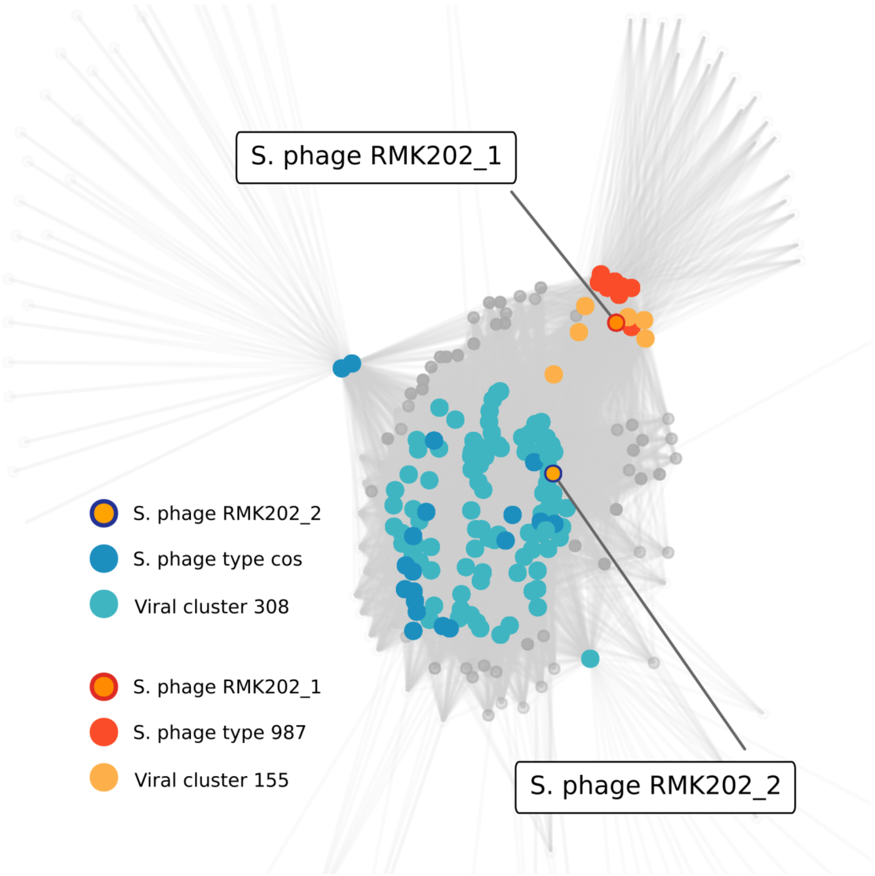


Figure 21. Similarity network of the two assembled *Streptococcus* phage genomes with previously sequenced phages and viral contigs based on vCONTACT v2.

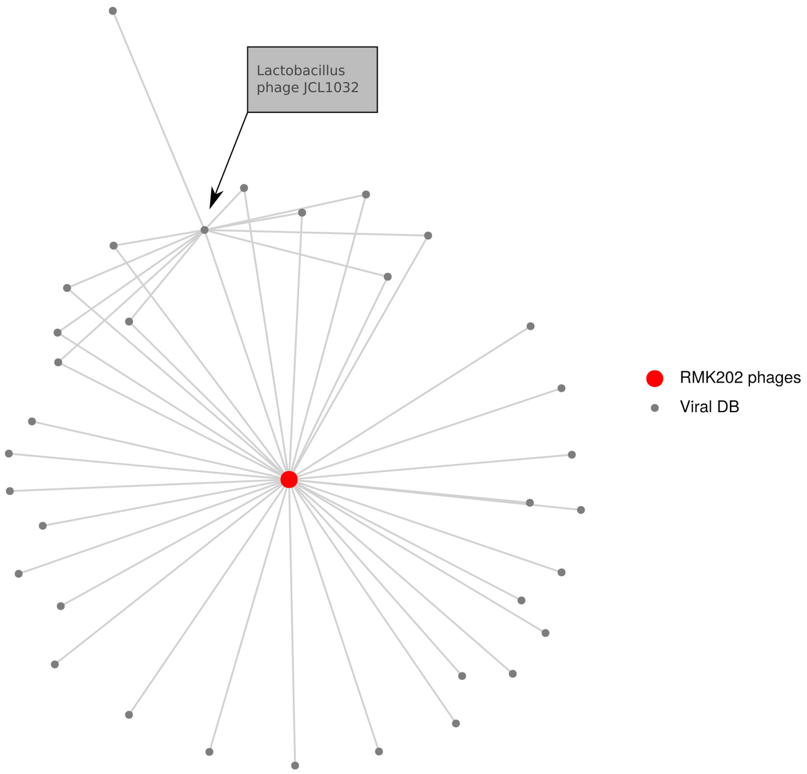


Figure 22. This plot will contain the *L. delbrueckii* phage network including the closest phage hit *Lactobacillus* phage JCL1032.

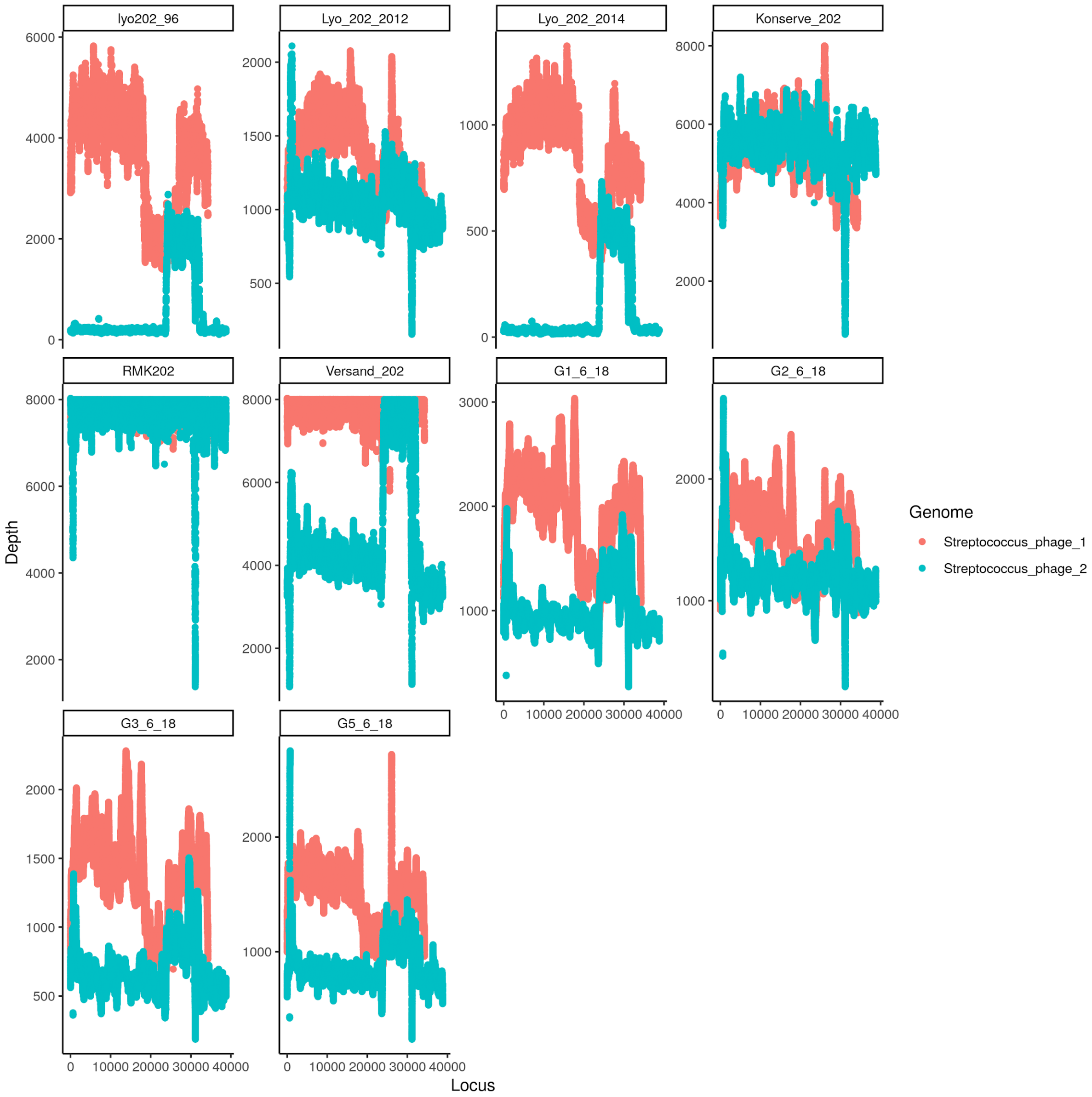


Figure 23. *Streptococcus* phage coverage in all metagenomic samples. The two genomes are very similar, especially in the lysis and lysogenic region, we thus see similar read recruitment by the different genomes. Note, the lysis & lysogenic region that is shared among the two phage genomes is recruiting the reads randomly to one or the other genome.

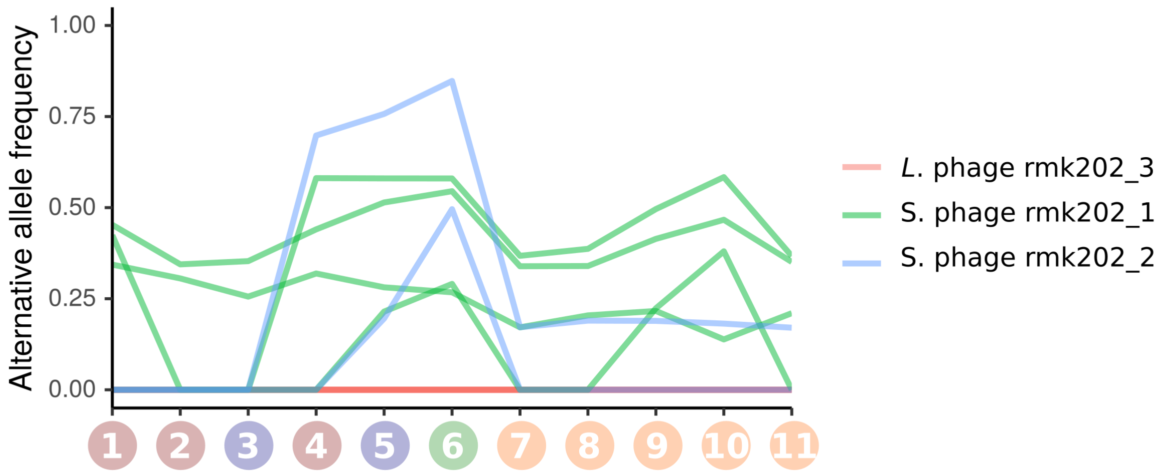


Figure 24. Alternative allele frequencies of the SNVs from the different phages. (The legend for the samples on the x-axis are illustrated in Fig. 2A).

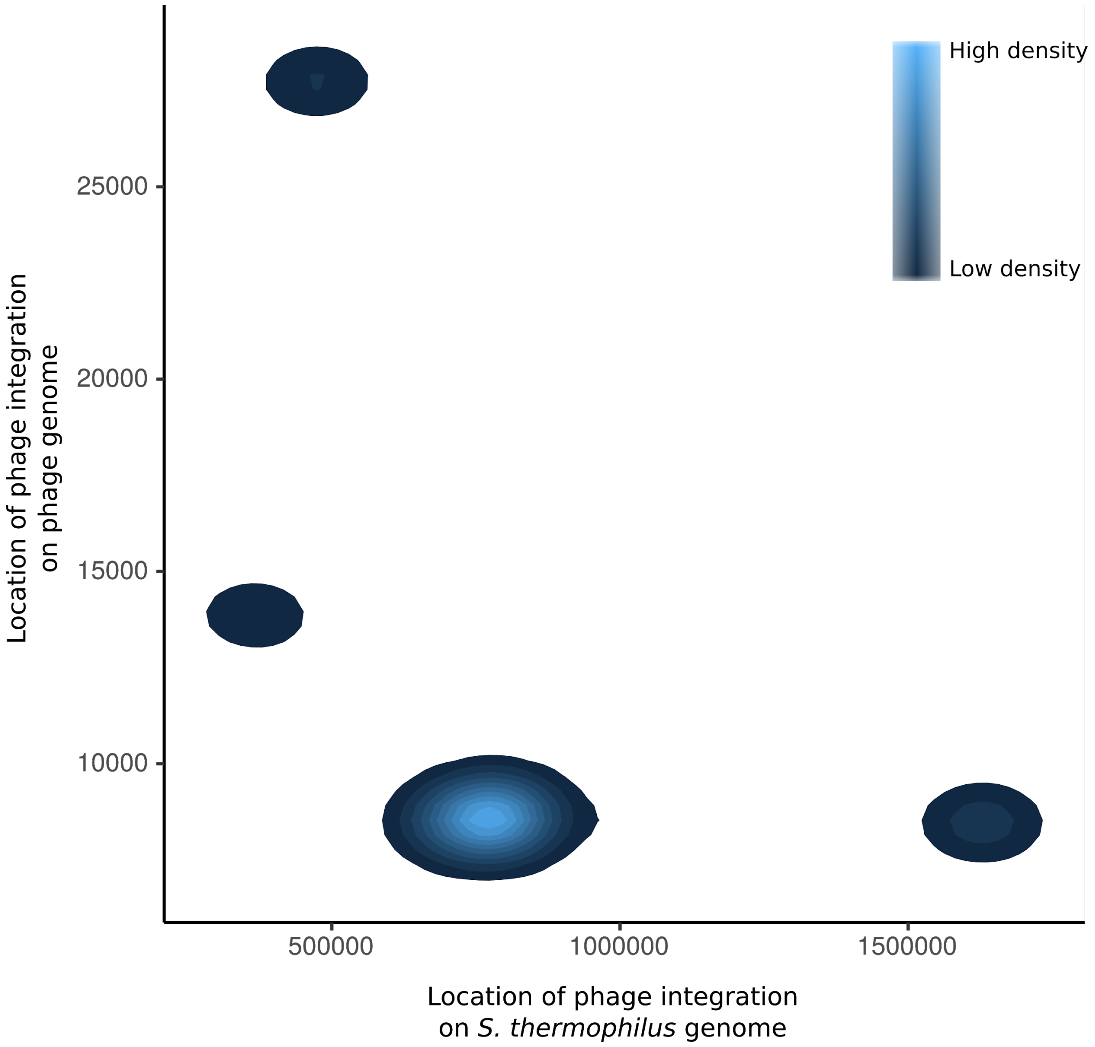


Figure 25. Four different putative integration sites were identified in the *S. thermophilus* (x-axis) and the phage genome (y-axis). The densities illustrate the number of reads mapping to the different locations.

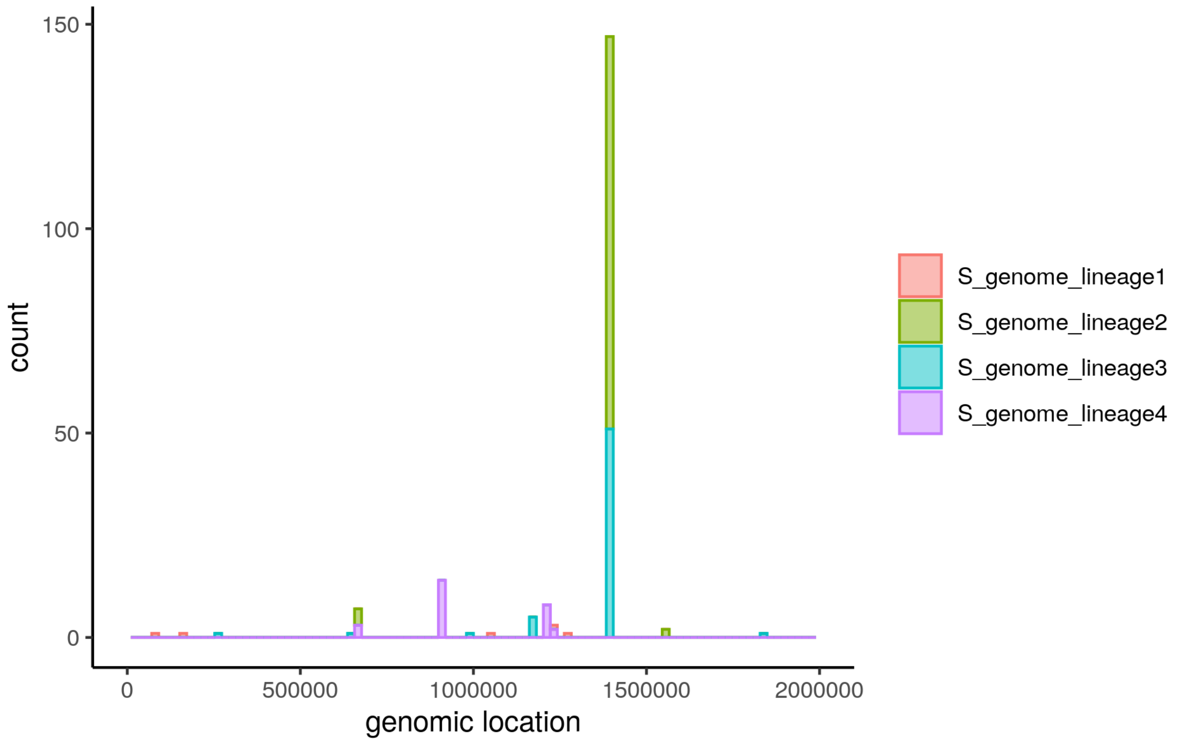


Figure 26. Genomic location on *S. thermophilus* where the lineage specific mate pair reads that span bacterial and phage genomes.

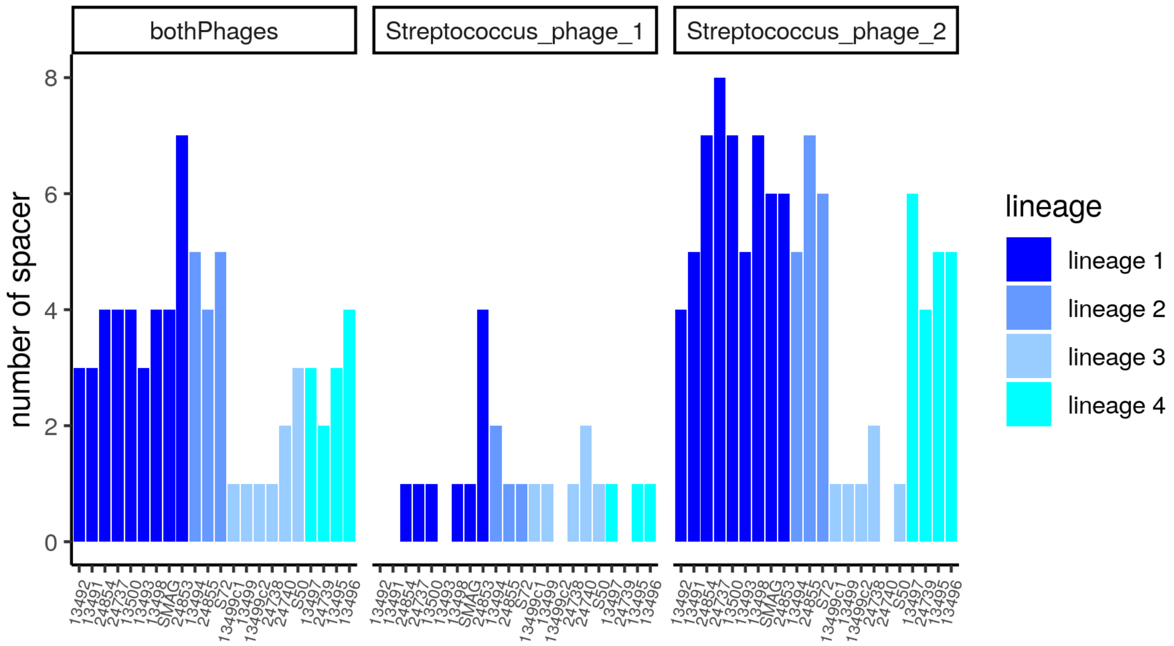


Figure 27. Number of CRISPR spacers that map to the two *Streptococcus* phages present in the starter culture. The color code for the genomes is illustrated in the legend and corresponds to the phylogenetic lineage.

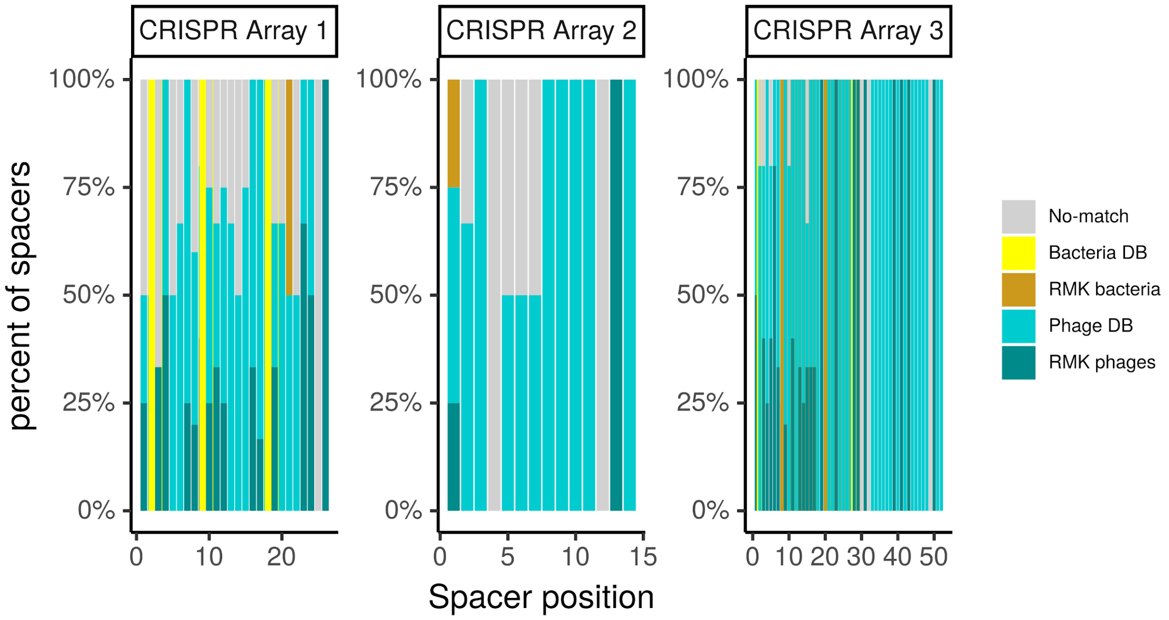


Figure 28. Origin of the CRISPR spacers plot according to the location on the CRISPR array.

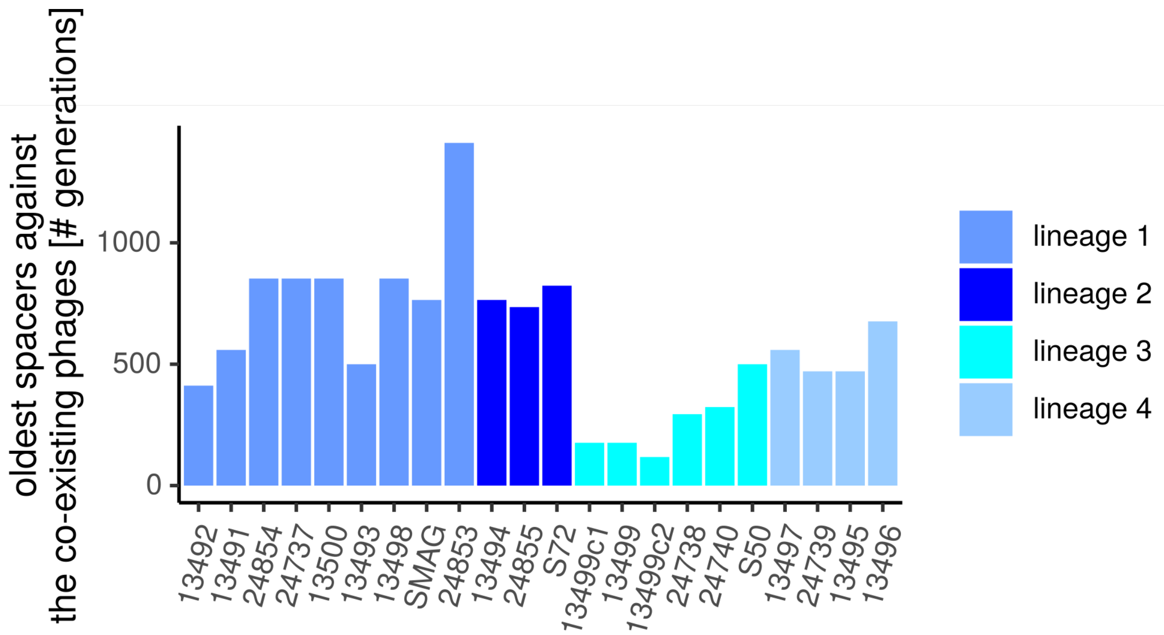

Figure 29. The estimated age (measured in generations) of the oldest spacers mapping against the co-existing phages for all *S. thermophilus* genomes. The number of generations was calculated by including the previously observed 0.024 spacer per generation turnover rate.

Table1: Statistics of all assembled or used genomes. GC stands for Guanine and Cytosine percentage in the genome. CRISPR stands for the number of CRISPR arrays annotated. BUSCO stands for the percent Busco completeness predicted.

| Species | Strain | Technology | # Contigs | Genome size | GC | Prophage | Plasmid | genes | Pseudogenes | rRNA | tRNA | transposase | CRISPR | BUSCO |
|---|---|---|---|---|---|---|---|---|---|---|---|---|---|---|
| *S. thermophilus* | 13494 | Nanopore&I. | 2 | 1882333 | 39 | 0 | 1 | 1946 | 230 | 88 | 67 | 79 | 3 | 99 |
| *S. thermophilus* | 13496 | Nanopore&I. | 1 | 1895875 | 39 | 0 | 0 | 1980 | 232 | 89 | 67 | 105 | 2 | 99 |
| *S. thermophilus* | 13498 | Nanopore&I. | 1 | 1865049 | 39 | 0 | 0 | 1934 | 223 | 88 | 66 | 81 | 3 | 99 |
| *S. thermophilus* | 13499 | Nanopore&I. | 1 | 1885735 | 39 | 0 | 0 | 1976 | 237 | 89 | 67 | 99 | 3 | 98 |
| *S. thermophilus* | 24737 | Nanopore&I. | 2 | 1868869 | 39 | 0 | 1 | 1938 | 229 | 91 | 68 | 81 | 3 | 99 |
| *S. thermophilus* | 24738 | Nanopore&I. | 1 | 1885982 | 39 | 0 | 0 | 1983 | 248 | 88 | 66 | 99 | 3 | 98 |
| *S. thermophilus* | 24739 | Nanopore&I. | 1 | 1934999 | 39 | 1 | 0 | 2027 | 234 | 89 | 67 | 103 | 2 | 99 |
| *S. thermophilus* | 24740 | Nanopore&I. | 1 | 1880058 | 39 | 0 | 0 | 1973 | 247 | 76 | 57 | 98 | 3 | 99 |
| *S. thermophilus* | 24853 | Nanopore&I. | 1 | 1847542 | 39 | 0 | 0 | 1920 | 221 | 89 | 67 | 73 | 3 | 99 |
| *S. thermophilus* | 24854 | Nanopore&I. | 2 | 1869650 | 39 | 0 | 1 | 1939 | 225 | 92 | 68 | 81 | 3 | 99 |
| *S. thermophilus* | 24855 | Nanopore&I. | 2 | 1926384 | 39 | 0 | 1 | 2010 | 261 | 110 | 85 | 85 | 3 | 99 |
| *S. thermophilus* | 13491 | Illumina | 46 | 1838081 | 39 | 0 | 0 | 1943 | 233 | 74 | 62 | 92 | 3 | 99 |
| *S. thermophilus* | 13492 | Illumina | 40 | 1835354 | 39 | 0 | 0 | 1923 | 229 | 59 | 50 | 89 | 3 | 99 |
| *S. thermophilus* | 13493 | Illumina | 54 | 1836485 | 39 | 0 | 0 | 1950 | 236 | 69 | 58 | 101 | 3 | 99 |
| *S. thermophilus* | 13495 | Illumina | 104 | 1803669 | 39 | 0 | 0 | 1937 | 251 | 59 | 50 | 91 | 2 | 95 |
| *S. thermophilus* | 13497 | Illumina | 63 | 1834883 | 39 | 0 | 0 | 1944 | 238 | 59 | 49 | 100 | 2 | 98 |
| *S. thermophilus* | 13499c1 | Illumina | 58 | 1838913 | 39 | 0 | 0 | 1960 | 248 | 63 | 53 | 100 | 3 | 99 |
| *S. thermophilus* | 13499c2 | Illumina | 60 | 1839354 | 39 | 0 | 0 | 1965 | 249 | 63 | 53 | 104 | 3 | 99 |
| *S. thermophilus* | 13500 | Illumina | 40 | 1831933 | 39 | 0 | 0 | 1938 | 230 | 74 | 62 | 87 | 3 | 99 |
| *S. thermophilus* | S50 | Illumina | 43 | 1842362 | 39 | 0 | 0 | 1921 | 231 | 47 | 42 | 74 | 3 | 98 |
| *S. thermophilus* | S72 | Illumina | 31 | 1835142 | 39 | 0 | 0 | 1897 | 225 | 49 | 43 | 68 | 3 | 99 |
| *S. thermophilus* | SMAG | MAG | 1 | 1879576 | 39 | 1 | 0 | 1975 | 227 | 89 | 67 | 75 | 3 | 99 |
| *S. thermophilus* | 19258 | Typestrain | 1 | 2102268 | 39 | 0 | 0 | 2230 | 287 | 75 | 56 | 100 | 2 | 99 |
| *L. delbrueckii* | 11141 | Illumina | 155 | 1994221 | 49 | 0 | 0 | 2112 | 191 | 96 | 80 | 199 | 3 | 97 |
| *L. delbrueckii* | 11142 | Illumina | 159 | 2017445 | 49 | 0 | 1 | 2140 | 170 | 96 | 79 | 215 | 2 | 98 |
| *L. delbrueckii* | 11143 | Illumina | 164 | 2002804 | 49 | 0 | 1 | 2142 | 185 | 91 | 73 | 225 | 3 | 97 |
| *L. delbrueckii* | 12104 | Illumina | 165 | 2003390 | 49 | 0 | 0 | 2168 | 197 | 100 | 80 | 247 | 3 | 97 |
| *L. delbrueckii* | 12105 | Illumina | 162 | 2012792 | 49 | 0 | 1 | 2170 | 196 | 99 | 81 | 241 | 3 | 98 |
| *L. delbrueckii* | 12107 | Illumina | 166 | 2007880 | 49 | 0 | 1 | 2129 | 180 | 104 | 82 | 202 | 3 | 98 |
| *L. delbrueckii* | 12109 | Illumina | 162 | 1985627 | 49 | 0 | 1 | 2079 | 187 | 88 | 70 | 171 | 3 | 98 |
| *L. delbrueckii* | 24776 | Illumina | 175 | 1954721 | 49 | 0 | 1 | 2041 | 172 | 98 | 78 | 126 | 3 | 97 |
| *L. delbrueckii* | 24777 | Illumina | 168 | 1933884 | 49 | 0 | 1 | 2011 | 166 | 94 | 77 | 120 | 3 | 97 |
| *L. delbrueckii* | 24778 | Illumina | 166 | 1910523 | 50 | 0 | 1 | 1920 | 153 | 62 | 54 | 48 | 3 | 97 |
| *L. delbrueckii* | 24779 | Illumina | 171 | 1955208 | 49 | 0 | 1 | 2032 | 164 | 91 | 74 | 125 | 3 | 97 |
| *L. delbrueckii* | 24780 | Illumina | 175 | 1953065 | 49 | 0 | 1 | 2035 | 170 | 92 | 74 | 131 | 3 | 97 |
| *L. delbrueckii* | 24781 | Illumina | 171 | 1943061 | 49 | 0 | 0 | 2021 | 163 | 92 | 74 | 126 | 3 | 97 |
| *L. delbrueckii* | 24782 | Illumina | 173 | 1888663 | 50 | 0 | 0 | 1974 | 162 | 95 | 77 | 126 | 1 | 97 |
| *L. delbrueckii* | 24783 | Illumina | 176 | 1946687 | 49 | 0 | 0 | 2020 | 167 | 94 | 77 | 122 | 3 | 97 |
| *L. delbrueckii* | 24798 | Illumina | 171 | 1956412 | 49 | 0 | 1 | 2032 | 172 | 92 | 77 | 127 | 3 | 97 |
| *L. delbrueckii* | LMAG | MAG | 1 | 2184491 | 49 | 0 | 1 | 2163 | 240 | 125 | 95 | 212 | 4 | 98 |
| *L. delbrueckii* | 20072 | Typestrain | 1 | 2165984 | 49 | 0 | 0 | 2152 | 228 | 124 | 94 | 210 | 1 | 98 |

### References


Fuchsmann, Pascal, Mireille Tena Stern, Patrick Bischoff, René Badertscher, Katharina Breme, and Barbara Walther. 2019. “Development and Performance Evaluation of a Novel Dynamic Headspace Vacuum Transfer ‘In Trap’ Extraction Method for Volatile Compounds and Comparison with Headspace Solid-Phase Microextraction and Headspace in-Tube Extraction.” *Journal of Chromatography A*. https://doi.org/10.1016/j.chroma.2019.05.016.

Vingataramin, Laurie, and Eric H. Frost. 2015. “A Single Protocol for Extraction of gDNA from Bacteria and Yeast.” *BioTechniques* 58 (3): 120–25.

Wick, Ryan R., Mark B. Schultz, Justin Zobel, and Kathryn E. Holt. 2015. “Bandage: Interactive Visualization of de Novo Genome Assemblies.” *Bioinformatics* 31 (20): 3350–52.

Borshchevskaya, L. N., T. L. Gordeeva, A. N. Kalinina, and S. P. Sineokii. 2016. “Spectrophotometric Determination of Lactic Acid.” Journal of Analytical Chemistry. https://doi.org/10.1134/s1061934816080037.

Danecek, P., A. Auton, G. Abecasis, C. A. Albers, E. Banks, M. A. DePristo, R. E. Handsaker, et al. 2011. “The Variant Call Format and VCFtools.” Bioinformatics. https://doi.org/10.1093/bioinformatics/btr330.

De Baets, Greet, Joost Van Durme, Joke Reumers, Sebastian Maurer-Stroh, Peter Vanhee, Joaquin Dopazo, Joost Schymkowitz, and Frederic Rousseau. 2012. “SNPeffect 4.0:

On-Line Prediction of Molecular and Structural Effects of Protein-Coding Variants." Nucleic Acids Research 40 (Database issue): D935–39.

Ellegaard, Kirsten M., and Philipp Engel. 2019. "Genomic Diversity Landscape of the Honey Bee Gut Microbiota." Nature Communications 10 (1): 446.

Garrison, Erik, and Gabor Marth. 2012. "Haplotype-Based Variant Detection from Short-Read Sequencing." arXiv.org, July. https://doi.org/arXiv:1207.3907.

Huerta-Cepas, Jaime, Damian Szklarczyk, Davide Heller, Ana Hernández-Plaza, Sofia K. Forslund, Helen Cook, Daniel R. Mende, et al. 2019. "eggNOG 5.0: A Hierarchical, Functionally and Phylogenetically Annotated Orthology Resource Based on 5090 Organisms and 2502 Viruses." Nucleic Acids Research 47 (D1): D309–14.

Jain, Chirag, Luis M. Rodriguez-R, Adam M. Phillippy, Konstantinos T. Konstantinidis, and Srinivas Aluru. 2018. "High Throughput ANI Analysis of 90K Prokaryotic Genomes Reveals Clear Species Boundaries." Nature Communications 9 (1): 5114.

Kolmogorov, Mikhail, Jeffrey Yuan, Yu Lin, and Pavel A. Pevzner. 2019. "Assembly of Long, Error-Prone Reads Using Repeat Graphs." Nature Biotechnology 37 (5): 540–46.

Krzywinski, Martin, Jacqueline Schein, Inanç Birol, Joseph Connors, Randy Gascoyne, Doug Horsman, Steven J. Jones, and Marco A. Marra. 2009. "Circos: An Information Aesthetic for Comparative Genomics." Genome Research 19 (9): 1639–45.

Li, Heng, and Richard Durbin. 2010. "Fast and Accurate Long-Read Alignment with Burrows-Wheeler Transform." Bioinformatics 26 (5): 589–95.

Martin, Marcel. 2011. "Cutadapt Removes Adapter Sequences from High-Throughput Sequencing Reads." EMBnet.journal 17 (1): 10–12.

Mikheenko, Alla, Vladislav Saveliev, and Alexey Gurevich. 2016. "MetaQUAST: Evaluation of Metagenome Assemblies." Bioinformatics 32 (7): 1088–90.

Milanese, Alessio, Daniel R. Mende, Lucas Paoli, Guillem Salazar, Hans-Joachim Ruscheweyh, Miguelangel Cuenca, Pascal Hingamp, et al. 2019. "Microbial Abundance, Activity and Population Genomic Profiling with mOTUs2." Nature Communications 10 (1): 1014.

Nurk, Sergey, Anton Bankevich, Dmitry Antipov, Alexey Gurevich, Anton Korobeynikov, Alla Lapidus, Andrey Prjibelsky, et al. 2013. "Assembling Genomes and Mini-Metagenomes from Highly Chimeric Reads." Lecture Notes in Computer Science. https://doi.org/10.1007/978-3-642-37195-0_13.

Shani, Noam, Dino Isolini, Daniel Marzohl, and Hélène Berthoud. 2021. "Evaluation of a New Culture Medium for the Enumeration and Isolation of Streptococcus Salivarius Subsp. Thermophilus from Cheese." Food Microbiology. https://doi.org/10.1016/j.fm.2020.103672.

Simão, Felipe A., Robert M. Waterhouse, Panagiotis Ioannidis, Evgenia V. Kriventseva, and Evgeny M. Zdobnov. 2015. "BUSCO: Assessing Genome Assembly and Annotation Completeness with Single-Copy Orthologs." Bioinformatics 31 (19): 3210–12.

Sjöqvist, C., L. F. Delgado Zambrano, J. Alneberg, and A. F. Andersson. n.d. "Revealing Ecologically Coherent Population Structure of Uncultivated Bacterioplankton with POGENOM." https://doi.org/10.1101/2020.03.25.999755.

Tatusova, Tatiana, Michael DiCuccio, Azat Badretdin, Vyacheslav Chetvernin, Eric P. Nawrocki, Leonid Zaslavsky, Alexandre Lomsadze, Kim D. Pruitt, Mark Borodovsky, and James Ostell. 2016. "NCBI Prokaryotic Genome Annotation Pipeline." Nucleic Acids Research. https://doi.org/10.1093/nar/gkw569.

Tonkin-Hill, Gerry, Neil MacAlasdair, Christopher Ruis, Aaron Weimann, Gal Horesh, John A. Lees, Rebecca A. Gladstone, et al. 2020. "Producing Polished Prokaryotic

Pangenomes with the Panaroo Pipeline.” Genome Biology 21 (1): 180.
Vaser, Robert, Ivan Sović, Niranjan Nagarajan, and Mile Šikić. 2017. “Fast and Accurate de Novo Genome Assembly from Long Uncorrected Reads.” Genome Research. https://doi.org/10.1101/gr.214270.116.
Vingataramin, Laurie, and Eric H. Frost. 2015. “A Single Protocol for Extraction of gDNA from Bacteria and Yeast.” BioTechniques 58 (3): 120–25.
